# Reversible conjugation of a CBASS nucleotide cyclase regulates immune response to phage infection

**DOI:** 10.1101/2023.09.29.560134

**Authors:** Larissa Krüger, Laura Gaskell-Mew, Shirley Graham, Sally Shirran, Robert Hertel, Malcolm F White

## Abstract

Antiviral defence systems build the prokaryotic immune system and their proper regulation is vital for survival and fitness. While it is important that they are readily available in case of infection, they need to be tightly controlled to prevent activation of an unnecessary cellular response. Here we describe how the bacterial cyclic oligonucleotide-based antiphage signalling system (CBASS) uses its intrinsic protein modification system to regulate the nucleotide cyclase. By integrating a Type II CBASS system from *Bacillus cereus* into the cognate host *Bacillus subtilis*, we show that the Cap2 protein conjugates the cyclase exclusively to the conserved phage shock protein A (PspA) in the absence of phage. This cyclase-PspA conjugation is reversed after infection by the isopeptidase Cap3. Finally, we propose a model in which the cyclase is held in an inactive state by conjugation to PspA in the absence of phage and that this conjugation is released upon infection, priming the cyclase for activation.

## Introduction

The evolution of bacteria and their viruses (phages) has been driven by the constant arms race between them. While bacteria developed more potent defence systems, phages evolved to overcome these systems. The best-known examples for anti-viral defence systems are the adaptive and innate immune system of eukaryotes and the restriction-modification and CRISPR systems of prokaryotes. While the eukaryotic systems evolved to a much higher complexity to function in multicellular organisms, they have their ancestral roots in the prokaryotic world. Thus, the eukaryotic cGAS enzyme of the cGAS-STING pathway is a homolog of the bacterial CD-NTase (cyclic nucleotidyl-transferase), which is part of CBASS (cyclic-oligonucleotide-based antiphage signalling system) ^1,2^. In fact, recent studies have revealed a large diversity of bacterial CD-NTase enzymes that are all related to cGAS and synthesize a diverse array of cyclic nucleotides in response to phage infection ^1–4^, initiating downstream effectors to modulate the bacterial immune response ^5–8^. In most cases this is thought to lead to dormancy or programmed cell death even though alternative strategies allowing bacterial survival have recently been described for other defence systems ^2,4,9–12^.

Although widespread in bacteria, key details of CBASS immunity remain unexplored. While it is a common feature of CRISPR systems to be activated by viral RNA/DNA, the signal that activates CD-NTases is more varied and in many cases still unknown. The eukaryotic homolog cGAS is stimulated by binding to double-stranded DNA or RNA in the cytosol, indicative of viral infection ^13–15^. Similarly, it has recently been observed that CD-NTases from Type I CBASS systems in *Staphylococcus schleiferi* bind viral RNA, thereby triggering cyclic dinucleotide synthesis ^16^. Phage capsid proteins have been implicated in triggering type II CBASS defence in *Pseudomonas aeruginosa* ^17^, while the CD-NTase DncV from *Vibrio cholerae* binds folate-like molecules and the disturbance of such during phage replication is thought to activate the enzyme ^18–20^. In Type III CBASS systems binding of certain phage peptides by HORMA-domain containing proteins changes their state and allows them to bind and activate the CD-NTase_21._

Type II CBASS operons contain genes encoding homologs of the ubiquitin ligation machinery and such systems account for 39 % of all CBASS systems, highlighting the relevance of the ancillary genes *cap2* and *cap3* ^4^. Very recently, two different groups have reported that the carboxy terminus of the CD-NTase is activated by ADP ribosylation and transiently linked via a thioester bond to the Cap2 enzyme, before it is finally conjugated to its target in the cell ^22,23^. While the first step of this posttranslational modification has been uncovered, the nature of the cellular target, the role of the regulation by conjugation of CD-NTase for immunity, as well as its link to phage infection remain largely unexplored.

To increase our understanding of CBASS regulation and to shed light onto the importance of the CD-NTase-conjugation, we studied the Type II CBASS system from the *Bacillus cereus* strain WPySW2. This CBASS operon consists of four genes encoding a cGAS/DncV-like nucleotidyltransferase, a ubiquitin activating enzyme E1 domain fused to an E2 ubiquitin transferase domain (Cap2), a JAMM (Jab1/MPN/Mov34 metalloenzyme) deubiquitinating enzyme (Cap3), and a Nuc-SAVED effector (Cap4) with a DNA endonuclease (Nuc) fused to a cyclic nucleotide binding SAVED domain (Fig. 1a). We transferred the system from *B. cereus* into the cognate host *Bacillus subtilis,* which allowed us to study this CBASS system *in vivo* and identify cellular components relevant for CBASS immunity. Using an unbiased cell extract separation approach we observe that the CD-NTase (henceforth “cyclase”) is in-part membrane associated and, in the presence of the Cap2 enzyme, conjugated to the phage shock protein A (PspA). PspA is the main effector of the phage shock protein (Psp) system, a universally conserved bacterial response to cell envelope stress resulting from phage infection, alkali shock, or membrane targeting antibiotics ^24^. On phage infection, Cap3-mediated deconjugation of the cyclase from PspA is observed. Based on our experimental data and published data on Psp systems, we hypothesize that CBASS uses the assembly of PspA in helical rods with an ESCRT-III-like fold as a platform to store individual cyclase monomers and inhibit their activity, hence preventing futile activation of the defence system.

**Figure 1.**
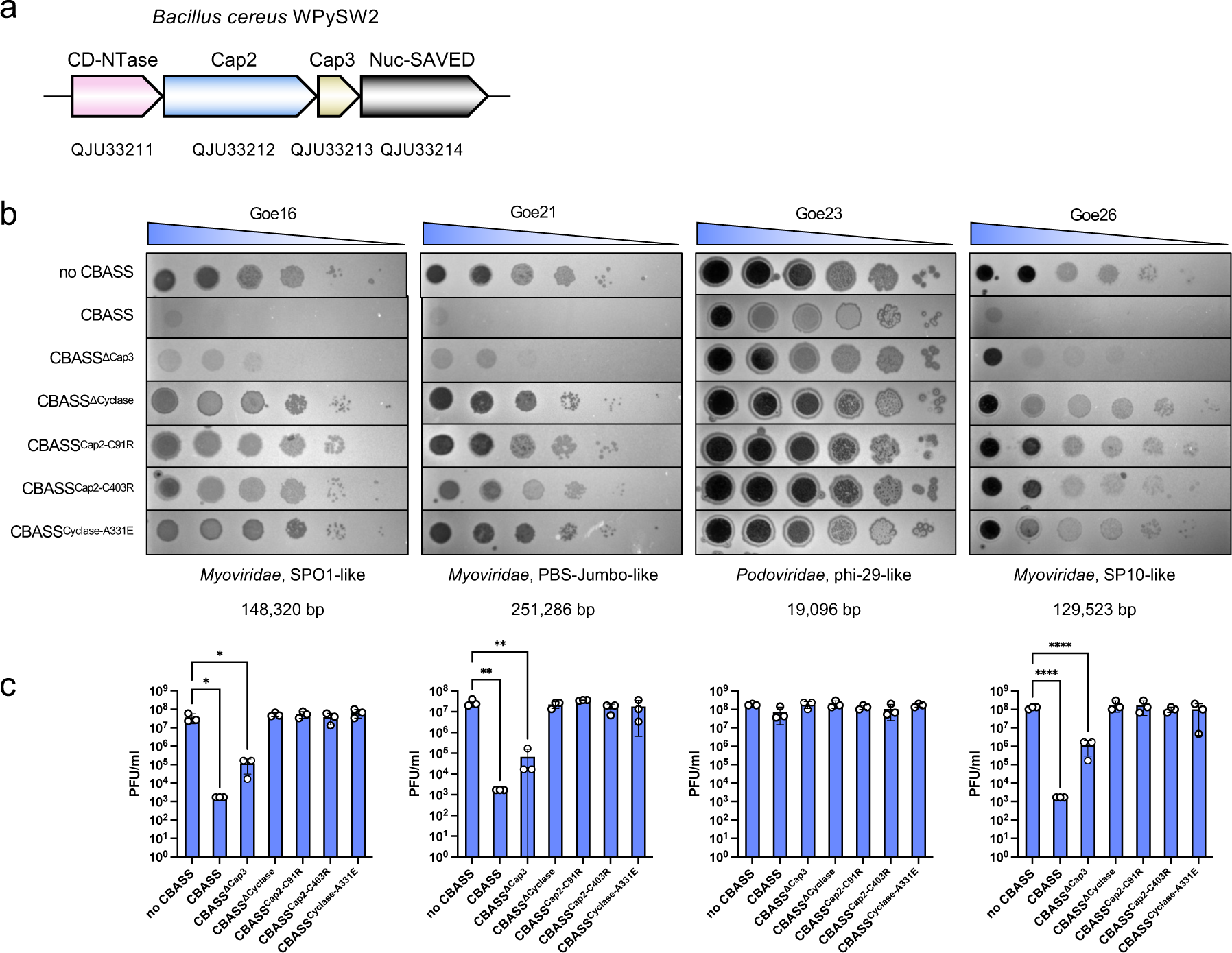
CBASS protects from phage infection. **a**, CBASS operon organisation in *Bacillus cereus* WPySW2 with GenBank accession numbers. **b**, Variants of the CBASS operon were integrated into the genome of *Bacillus subtilis* Δ6. Exponentially growing cells were infected with dilutions of the respective phage. Representative plates of three biological replicates are shown. The family, the genus, and the genome size of each phage are depicted below the plaque assays. **c**, Plaque forming units (PFU/ml) of infection drop assays in b. Data are presented as mean values ± SD. Statistical analysis was performed using a one-way ANOVA, followed by Dunnett’s multiple comparisons test (*P < 0.0129, **P < 0.0022, ****P < 0.0001).

## Results

### The *B. cereus* CBASS system is functional in *B. subtilis*

In order to study the CBASS system of *B. cereus* WPysSW2 (*Bce*) *in vivo*, the operon (Fig. 1a) was introduced into the genome of *B. subtilis* Δ6 and expressed under a strong, constitutive *B. subtilis* promoter (PdegQ) ^25,26^. *B. subtilis* Δ6 was used for the infection experiments because it is deficient of the two prophages (SPβ and PBSX) and the three prophage-like elements (prophage 1, prophage 3, and skin) in the parental *B. subtilis* 168 strain ^27^. The protective effect of CBASS against phage infection was tested by infecting the *B. subtilis* Δ6 parental strain (from now on referred to as “no CBASS”), as well as strains containing either the full or partial CBASS operon, with four novel *B. subtilis* phages. We compared the genomes of the four phages to known *B. subtilis* phages and classified them accordingly (Extended Data Fig. 1a). Therefore, Goe16, Goe21, and Goe26 were classified to belong to the *Myoviridae* family and the SPO1-like phages, PBS Jumbophage and SP10-like virus genus, respectively. Goe23 is a representative of the *Podoviridae* family and the Phi29-like genus (for more details see Supplementary Table 1).

We observed that the type II CBASS system was able to protect bacteria from infection with all four phages (Figure 1b,c, Extended Data Fig. 2). CBASS conferred 10^4^-fold defence compared to the parental strain for phages Goe16, Goe21 and Goe26. We did not observe a reduction in PFU count for Goe23 (Fig. 1b,c), however, the morphology of the plaques changed from a clear to a much more turbid plaque with no clear centre of lysis (Fig. 1b). Additionally, we did observe protection in liquid medium supporting the response of CBASS to Goe23 (Extended Data Fig. 2a). Thus, the CBASS*^Bce^* system is able to confer immunity against four different classes of phages with very different characteristics (Fig. 1b). Goe21 is a jumbo phage, a class of phage known to shield their DNA in a nucleus-like structure. Phage Goe16 and Goe26 presumably use modified bases in their DNA. Goe16 encodes for homologous enzymes involved in the synthesis of hydroxymethyluracil nucleotides in phage SPO1 ^28^. Goe26 encodes for homologs of enzymes supposedly capable of hypermodification of thymines in SP10 as suggested by domain conservation ^29^. These flexibilities in both viral DNA storage and composition could be an indicator that viral DNA is not the indicator of infection detected by CBASS*^Bce^*. Likewise, direct detection of specific phage proteins seems unlikely given the diversity of phages studied here.

In order to assess the essential components of CBASS, we deleted parts of the operon, or introduced point mutations in relevant residues to generate protein variants. The double deletion of the *cap2* and *cap3* genes led to a complete loss of immunity, similar to the effect of deletion of the *cyclase* gene. In contrast, the absence of Cap3 (the JAMM protease) alone reduced, but did not abolish, CBASS immunity (Fig. 1b, c). This indicates that the potential deconjugating activity of Cap3 is not essential, but is important, in the CBASS*^Bce^* system. In contrast to Cap3, Cap2 was essential for CBASS immunity as mutation of the catalytic residues of either the E2 or E1 domain, led to loss of immunity (Fig. 1b, c).

The C-terminal tails of CD-NTases are highly conserved even among distantly related Type II CBASS systems and the last amino acid of CBASS cyclases is in most cases an Ala or Gly residue^22^. This C-terminal residue is the site of conjugation by Cap2 to a lysine residue on cellular targets ^22^. To investigate the importance of the C-terminal Ala in our system, we mutated Ala331 to a negatively charged Glu residue (A331E) and observed that immunity was lost (Fig. 1bc).

### The *Bce* cyclase is conjugated to Phage Shock Protein A

Eukaryotic cGAS associates with the plasma membrane by selective interaction of its N-terminal domain with phosphoinositide, a feature that is important for the regulation of the enzyme ^30^. The subcellular localisation of the bacterial CD-NTase enzymes could be an important aspect of their regulation, but has not yet been explored. We addressed this by overexpressing the *Bce* cyclase in *B. subtilis* strains containing genomic integrations of Cap2 and the effector. We separated the cytosolic from membrane fractions and purified the cyclase by affinity chromotography via protein A-coupled beads saturated with cyclase-specific antibodies. We observed that, while most of the overexpressed cyclase was cytosolic, there was a substantial amount of the protein in the membrane fractions (Fig. 2a, Extended Data Fig. 3a). We observed this for the full-length cyclase as well as a truncated mutant, in which the flexible C-terminal tail had been deleted.

**Figure 2.**
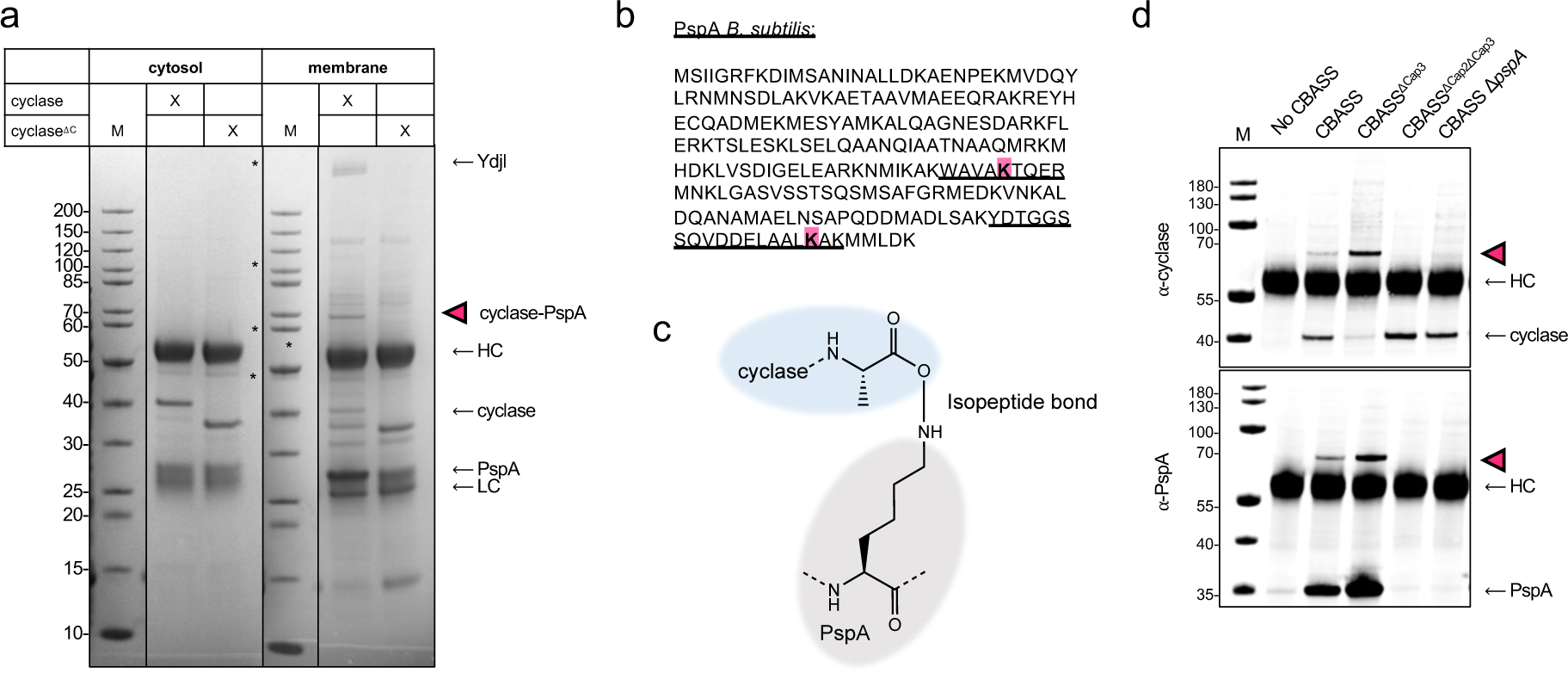
The cyclase partitions between cytoplasm and membranes and is conjugated to the phage shock protein A (PspA) *in vivo*. **a**, The full-length or truncated (1-300) cyclase were overexpressed in *B. subtilis* in the absence of Cap3 (strain LK40). Cultures were harvested at exponential growth phase and cell lysates separated into cytosolic and membrane fractions. The cyclase was purified from both fractions using protein A coupled beads and a cyclase specific antibody. Fractions of the purified cyclase were run on SDS-PAGE. The gel shows the elution fractions from the purification. In the membrane fraction additional higher molecular weight bands can be detected for the full-length cyclase with a prominent band below the 70 kDa band (magenta arrow). Abbreviations: HC: heavy chain; LC: light chain; ΔC: deletion of C-terminus of cyclase (aa 302-331). **b**, Amino acid sequence of PspA from *B. subtilis* with the modified lysine residues highlighted (magenta) and the respective tryptic peptides underlined as they were identified with mass spectrometry analysis from the gel in a. **c**, Depiction of the conjugation site between PspA(K147/K220) and the cyclase (A331). **d**, The cyclase was purified from *B. subtilis* cell extracts expressing genomically-integrated versions of CBASS (strains LK18 (control), LK06 (CBASS), LK20 (CBASS^ΔCap3^), LK10 (CBASS^ΔCap2ΔCap3^), LK51 (CBASS Δ*pspA*)), separated by SDS-PAGE and analyzed by Western blot with specific antibodies (⍺-cyclase, ⍺-PspA). All experiments were conducted with at least two biological replicates.

In the membrane fractions, we observed a number of additional bands (at 28, 65 and >200 kDa) associated with the full-length cyclase (Fig. 2a, Extended Data Fig. 3a). Analysis by mass spectrometry showed the 65 kDa band (magenta arrow) contained almost equal amounts of the cyclase and a protein identified as phage shock protein A (PspA), suggestive of a 1:1 conjugated species (Extended Data Fig. 3ab).

In *B. subtilis*, the *pspA-ydjG-ydjH-ydjJ* operon is under control of the alternative extracytoplasmic function sigma factor σW which regulates expression of several genes in response to alkali stress and infection with phage SPP1 ^31^. In the higher molecular weight band (>200 kDa) we identified peptides of the YdjI protein (Fig. 2a; Extended Data Fig. 3b). YdjI is a stomatin, prohibitin, flotillin, and HflK/C-domain protein which localizes into liquid disordered membrane regions, and homologs assemble into higher oligomeric structures ^32–34^. Interestingly, together with the two membrane integral proteins YdjG and YdjH, it is essential for membrane localisation of PspA in *B. subtilis* ^32,35^. The prominent band at 25 kDa proved to contain the PspA protein. Comparing the intensity of the band in the SDS gel of both the PspA monomer and the cyclase, it became apparent that PspA was present in high molar excess (Extended Data Fig. 3b). It has been described that PspA proteins assemble in helical rods with an ESCRT-III-like fold ^36,37^, and such PspA foci have been observed in *B. subtilis* microscopically^32^. Thus, we hypothesize that the cyclase associates with and is conjugated to PspA rods *in vivo*.

To confirm that the cyclase and PspA proteins were indeed conjugated by an isopeptide bond, and identify the sites on each protein, we searched the MS data for PspA peptides carrying the last tryptic peptide of the cyclase, KPGGFA, which results in a predicted mass increase of 557.3 Da. We observed this mass shift on two residues, K147 and K220, of PspA (Fig. 2b, Extended Data Fig. 3b). This indicates that the C-terminal carboxyl group of the cyclase forms an isopeptide bond with the ε-amine of a lysine residue in PspA, reminiscent of ubiquitination in eukaryotes (Fig. 2c).

As our previous experiments were performed in the absence of Cap3, as well as with overexpression of the cyclase to increase our chances of identifying the target of cyclase-conjugation, we were now interested in which components of CBASS were required for cyclase conjugation under more realistic conditions *in vivo*. For this purpose, we purified cyclase from *B. subtilis* cells expressing genomically-integrated CBASS and performed Western Blot analysis to determine the level of conjugation in the different strains. As expected, the cyclase-PspA conjugate was only formed in the presence of cyclase, Cap2 and PspA, indicating that Cap2 is indeed the enzyme responsible for cyclase ligation to PspA (Fig. 2d). Interestingly, in the absence of PspA, we did not observe conjugation to an alternative target (Fig. 2d). Monomeric PspA clearly co-purified with the cyclase-PspA conjugate in high molar excess, providing further evidence that the conjugate is part of large PspA assemblies (Fig. 2d). The fact that the presence of Cap3 prevents formation of the cyclase-PspA-conjugate strongly indicates that Cap3 functions as the isopeptidase cleaving the cyclase-PspA-conjugate *in vivo*, which is in agreement with previous observations ^22,38^. We then separated the cell extract of the same strains into membrane and cytosolic fractions and purified the cyclase from the cytosolic fraction to concentrate the sample to the same volume as the membrane fraction. Western blot analysis showed that the cyclase partitions between both fractions (Extended Data Fig. 4). The association with the membrane fractions was independent of Cap2, Cap3 and PspA, providing evidence that the cyclase itself could interact with membranes (Extended Data Fig. 4).

### Conjugation *in vitro*

Knowing the full-length cyclase, Cap2 and PspA are required for conjugation *in vivo*, we aimed to reconstitute conjugation *in vitro*. For this purpose, we incubated the purified proteins in our conjugation assay and analysed the reaction mixtures by SDS-PAGE and Western blot using antibodies raised against the cyclase and PspA. With this, we observed conjugation of the cyclase to PspA in the presence of Cap2 and ATP (Fig. 3a). In addition to conjugation to PspA, the cyclase formed higher MW species in the presence of Cap2 and ATP which were stable in the presence of the reducing agent DTT, indicative of Cap2-cyclase thioester-intermediates of the conjugation reaction, as observed previously by others ^22,23^. Indeed, analysis of the respective higher MW species by mass spectrometry confirmed the presence of cyclase and Cap2 peptides in these bands (Extended Data Fig. 5a). The physiological relevance of these species is unclear, as they were not observed in *B. subtilis* (Fig. 2d).

**Figure 3.**
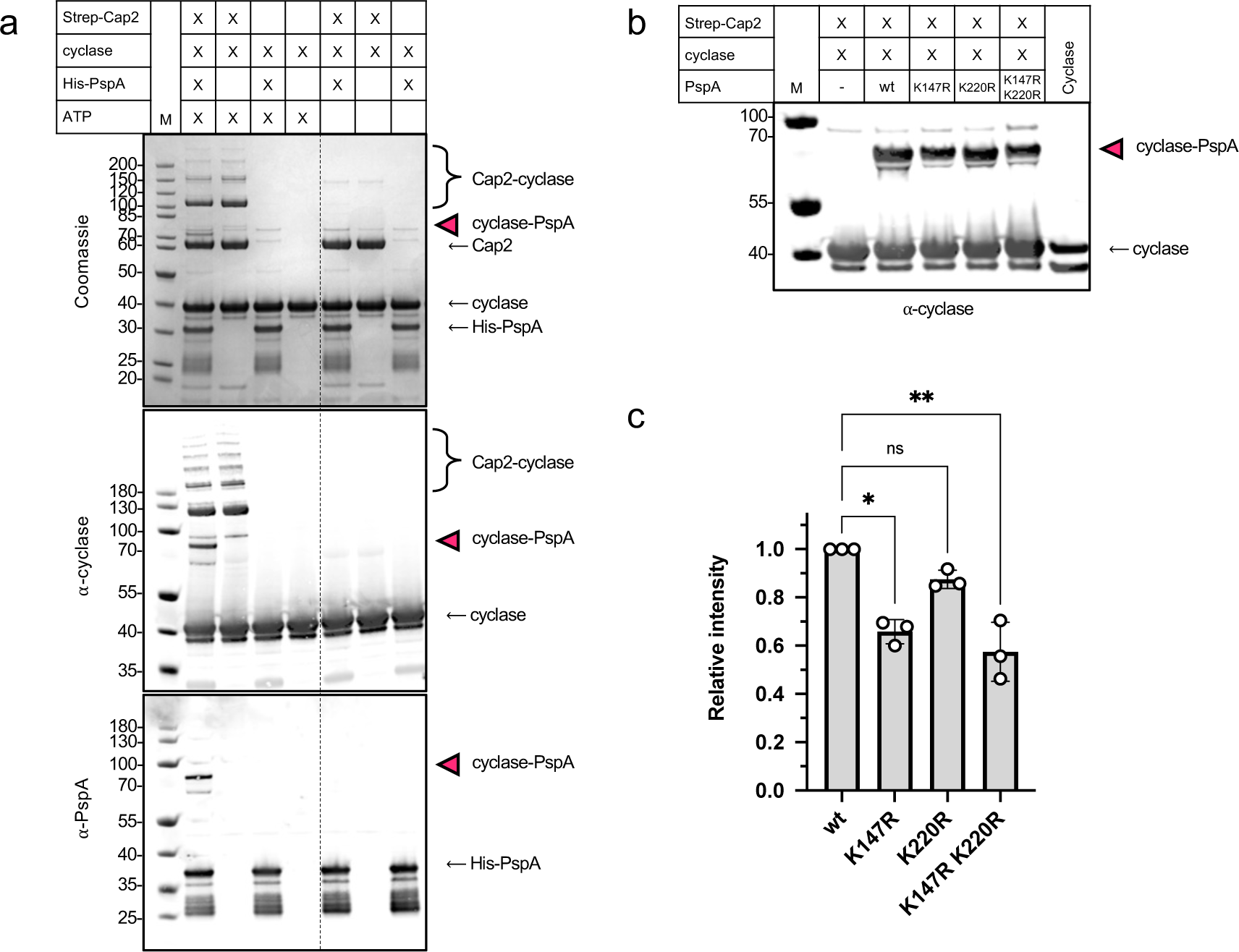
Cap2 conjugates the cyclase to PspA. **a**, *In vitro* conjugation of the cyclase to PspA. Reaction mixture contained 5 µM Strep-Cap2, 5 µM cyclase and 10 µM PspA, 10 mM MgCl_2_, 1 mM ATP, 100 mM Tris-HCl pH 8, 150 mM NaCl. Samples were incubated for 30 min at 25 °C, separated by SDS-PAGE and analysed by Western blot with specific antibodies (⍺-cyclase, ⍺-PspA). The cyclase-PspA conjugate is marked with magenta arrows. Strep-Cap2: 67.8 kDa; cyclase: 37.8 kDa; cyclase^ΔC^: 34.2 kDa; 6xHis-V5-PspA: 30.2 kDa. **b**, *in vitro* conjugation with PspA lysine mutants (reaction mix as in a). **c,** Quantification of b. Data are presented as mean values ± SD. Statistical analysis was performed using a one-way ANOVA, followed by Dunnett’s multiple comparisons test (**P = 0.0037; *P = 0.0151). All experiment were conducted with three biological replicates.

Having identified potential modification sites on PspA, we were interested in whether the conjugation to PspA was exclusive to these lysine residues. Thus, we created variant PspA proteins in which we exchanged either K147 or K220 for an arginine residue. We then tested the variants in our *in vitro* conjugation assay, observing that the K147R single mutant showed a 35 % reduction in conjugation, while the K220R mutant was still very efficiently targeted for conjugation (Fig. 3bc, Extended Data Fig. 5b). We then combined the K147R and K220R mutations and tested the double mutant in the conjugation assay, observing that the additional mutation of K220 in the K147R mutant led only to a minor reduction in conjugation (Figure 3bc, Extended Data Fig. 5b). This indicates that K147 is a preferred residue for conjugation, while there is a certain flexibility regarding lysine residues on PspA targeted for conjugation. The low sequence conservation between PspA homologs may explain why Cap2 requires flexibility in target recognition (Extended Data Fig. 5c). Based on this we hypothesize that interaction between Cap2 and PspA initiates conjugation of the C-terminus of the cyclase to an exposed lysine residue on the outward-facing C-terminus of PspA assembled in the large helical filament.

### Relevance of conjugation *in vivo*

To understand the relevance of conjugation during phage infection, we infected cells expressing either the full CBASS system or CBASS^ΔCap3^ with phage Goe23. The phage Goe23 belongs to the very well characterized phi29-like family and has a minimal genome. It also showed the most clear plaques (Fig. 1b), supposedly inducing the most drastic CBASS response, and, thus, making it an ideal candidate to study our system *in vivo*. Cells were harvested before- and 5, 15, and 30 min post-infection, and the membranes purified. In the Western blots we observed that infection did not increase formation of the conjugate at any time point. In contrast, we observed a Cap3-dependent decrease of the cyclase-PspA conjugate 30 min after infection (Fig. 4a,b). These data are consistent with a model involving an inhibitory effect of conjugation on the cyclase which is released upon infection with phage. To further evaluate this, we deleted *pspA* from the genome and infected the bacteria with each of the four phages. Deletion of *pspA* had no negative effect on immunity, suggesting that conjugation to PspA is not essential for the eventual activation of the cyclase (Fig. 4c).

**Figure 4.**
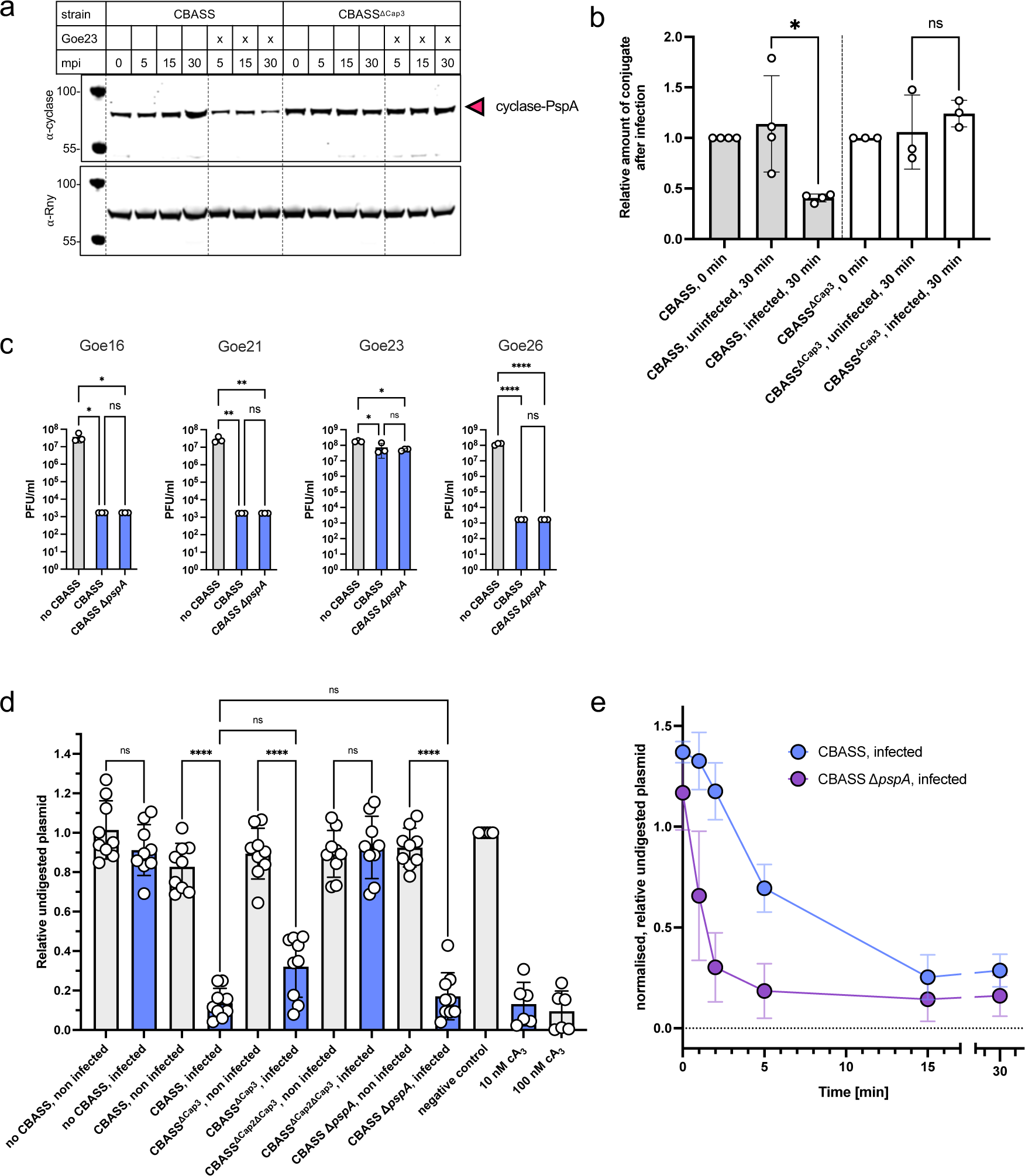
Cyclase conjugation to PspA responds to phage infection. **a**, Strains harboring genomic integration of the CBASS operon (strains LK06 (CBASS), LK20 (CBASS^ΔCap3^)) were grown until exponential growth phase and, if indicated, infected with phage Goe23 to an MOI of 2. Cultures were grown for an additional 5, 15, or 30 min before harvesting. Cell lysates were separated into cytosolic and membrane fractions. Membrane fractions were adjusted in protein content, separated by SDS-PAGE, and the conjugate was detected by Western blot with a cyclase-specific antibody. A specific antibody raised against the RNase Y protein served as a loading control. Abbreviations: mpi: minutes post infection **b**, Quantification of reduction of cyclase-PspA conjugate upon phage infection. Data is shown relative to wild type CBASS at time point 0 min. The experiment was conducted with n = 3/ n = 4 biologically independent samples. Data are presented as mean values ± SD. Statistical analysis was performed using a one-way ANOVA, followed by Tukey’s multiple comparisons test (*P=0.0129). **c**, Infection assay of *B. subtilis* strains containing CBASS components as indicated. Plaque forming units (PFU/ml) of infection drop assays were calculated from three biological replicates. No CBASS and CBASS data has been replotted from Fig. 1c. **d**,. Nucleotides were extracted from infected (Goe23, MOI=2) or uninfected *B. subtilis* cells carrying CBASS versions as indicated. These nucleotides and synthetic cA_3_ were tested for their ability to activate the CBASS effector Nuc-SAVED in a plasmid cleavage assay. **e**, quantification of uncleaved plasmid after 30 min incubation of the nucleotide extracts with the Nuc-SAVED protein. The experiment was conducted with three biological replicates with three technical replicates each. Data are presented as mean values ± SD. Statistical analysis was performed using a one-way ANOVA, followed by Tukey’s multiple comparisons test (****P<0.0001).

To tackle the suspected inhibitory effect of the cyclase conjugation further, we grew *B. subtilis* cultures expressing CBASS and infected the cultures with phage Goe23. These cultures were harvested 0, 15 and 30 min after infection and the intracellular nucleotides were extracted. We proceeded to test their ability to activate the cognate CBASS Nuc-SAVED effector in a plasmid cleavage assay. We quantified the amount of uncleaved plasmid after incubation of Nuc-SAVED with the nucleotide extracts or synthetic (3’, 3’, 3’) cA_3_ and observed plasmid degradation only 30 min after infection, but not at time 0 or 15 min after infection (Extended Data Fig. 6). This suggests that the cyclase is inactive in the absence of phage and that CBASS responds rather late during the infection cycle, which is in agreement with previous observations by others ^3^. To further investigate cyclase activity, we repeated the experiment with different CBASS mutant strains and observed that the second messenger nucleotide could only be detected in extracts from cells expressing a functional CBASS system (Fig. 4d, Extended Data Fig. 7, Fig. 1b,c). The only mutant strain that did not produce the second messenger after phage infection was the CBASS^ΔCap2ΔCap3^ strain that is unable to confer immunity (Fig. 4d, Extended Data Fig. 7, Fig. 1b,c). From this we can conclude that the cyclase is inactive in the absence of phage and that Cap2 is essential for activation. As we did not observe a difference in Nuc-SAVED cleavage activity with nucleotides extracted from CBASS or CBASS Δ*pspA* in the initial assay, we decided to measure Nuc-SAVED activity over a broader range of time. With this we observed earlier activation of Nuc-SAVED when incubated with extracts originating from CBASS Δ*pspA* compared with those from bacteria encoding PspA (Fig. 4e, Extended Data Fig. 8), indicating increased second messenger production in the absence of PspA at early time points. This supports the idea that the cyclase needs to be deconjugated prior to activation and that the initial response is faster in the *pspA* deletion mutant. This slow deconjugation step may in part explain the delayed kinetics of CBASS defence *in vivo* ^3^. This effect may be more pronounced in the native host where CBASS expression might generally be much lower and the ratio of conjugated/unconjugated cyclase is more drastically affected upon infection compared to our system.

### Conjugation in the heterologous *E. coli* host

As it has been previously observed that cyclases from *V. cholerae*, *E. coli* and *P. aeruginosa* can form Cap2-dependent higher MW species when expressed in *E. coli* ^22,38^, we co-expressed His-tagged cyclase and Cap2 in *E. coli* and purified the cyclase by immobilized metal affinity chromatography. In contrast to previously reported observations ^22,38^, few higher MW species were observed, suggesting that the final target for conjugation was missing in our heterologous system (Fig. 5a, lane 5). Thus, we co-expressed His-tagged cyclase and Cap2 together with PspA in *E. coli* and observed the formation of higher MW bands in the SDS gel (Fig. 5a, lane 1). We analysed these bands by mass spectrometry, and they did contain peptides of both the cyclase and the PspA protein, showing that almost half of the cyclase was conjugated to PspA (Extended Data Fig. 9a). Moreover, we identified K147 to be the major target for modification by the C-terminus of the cyclase, confirming our initial results (Extended Data Fig. 9a, Fig. 2b). In agreement with our experiments in *B. subtilis* (Fig. 2a), the cyclase-PspA conjugate was not formed with the truncated cyclase (Fig. 5a, lane 4). Conjugation was also abolished when the cyclase and PspA were co-expressed with a catalytic dead mutant of Cap2 (Fig. 5a, lane 2/3). These data provide strong support for the hypothesis that PspA is the relevant acceptor protein for conjugation with the cyclase.

**Figure 5.**
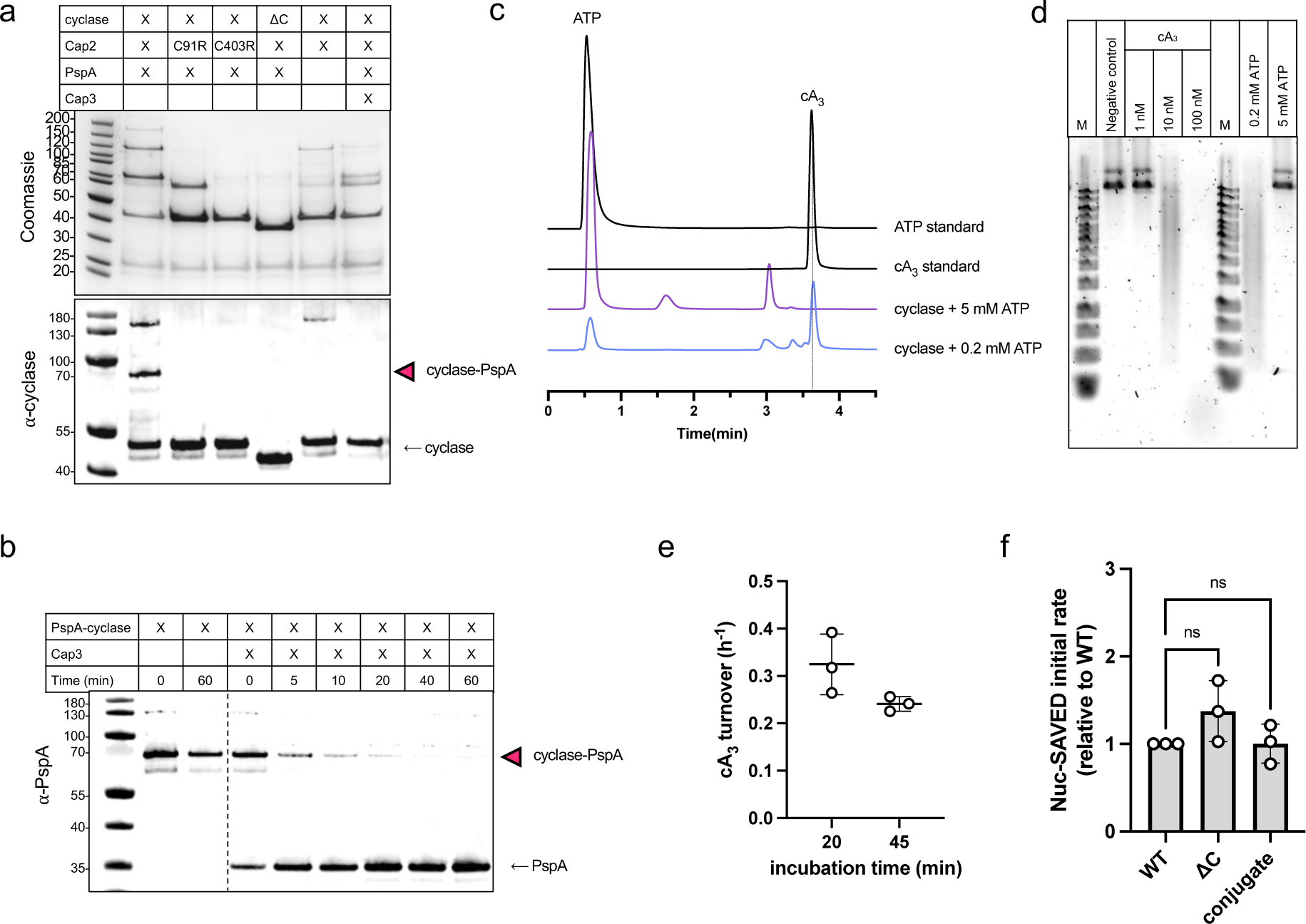
Conjugation in the heterologous *E. coli* host. **a**, Co-expression of the respective CBASS components in *E. coli* and purification via His-tagged cyclase. Separation by SDS-PAGE and the cyclase and conjugate were detected by Western blot with a cyclase-specific antibody. The presence of PspA peptides in the conjugate band was confirmed by mass spectrometry. Abbreviations: ΔC: deletion of C-terminus (aa 302-331). **b**, the cyclase-PspA conjugate was incubated together with the Cap3 ortholog from *Cytobacillus oceanisediminis* and the cleavage was followed by Western blot with a PspA-specific antibody (see also Extended Data Fig. 9c). **c**, reaction products from a cyclase reaction (50 µM cyclase, 0.2/ 5 mM ATP, 45 min incubation at 37 °C) were analysed by HPLC. The y-axis is rescaled for presentation. The experiment was conducted with three replicates and a representative trace is shown. See Extended Data Figure 10 for replicates. **d**, The reaction products from c were tested for their ability to activate the CBASS effector Nuc-SAVED in a plasmid cleavage assay. The experiment was conducted with three replicates and a representative gel is shown. **e**, the turnover number of the cyclase with respect to cA_3_ production with 0.2 mM ATP as substrate was calculated from the HPLC data by measuring the area under the peak and fitting the data to a cA_3_ standard curve (see Extended Data Fig. 10 for HPLC traces of replicates and linear regression of cA_3_ standard). Data are presented as mean values ± SD. **f,** the activity of the wild type cyclase, the truncated cyclase (1-301) and the cyclase-PspA conjugate was measured in a fluorescent DNA cleavage assay with the effector protein Nuc-SAVED. The initial rate of Nuc-SAVED was calculated and normalized to the protein amount and set relative to the wild type protein. The experiment was conducted with three replicates. Data are presented as mean values ± SD. Statistical analysis was performed using a one-way ANOVA, followed by Dunnett’s multiple comparisons test (ns = 0.1679).

As expected from previous experiments, co-expression with Cap3 drastically reduced the formation of the cyclase-PspA conjugate (Fig. 5a, lane 6). Unfortunately, we were unable to express the *Bce* Cap3 protein and used instead a Cap3 ortholog from *Cytobacillus oceanisediminis* (61 % sequence identity to Cap3*^Bce^*) to confirm the activity of Cap3 *in vitro*. To obtain the substrate, we purified the cyclase from the co-expression and separated monomeric cyclase from the cyclase-PspA conjugate by size exclusion chromatography (Extended Data Fig. 9b). Incubation of the purified cyclase-PspA conjugate with Cap3*^Coc^* showed that Cap3 is indeed the isopeptidase cleaving the conjugate (Fig. 5b, Extended Data Fig. 9c). We compared the mass of non-conjugated cyclase to deconjugated cyclase by intact mass spectrometry and determined that they had the same mass (Extended Data Fig. 9e). Additionaly, we treated the two proteins with the peptidase AspN and analysed the peptides by mass spectrometry, identifying the C-terminal peptide of the cyclase in both versions, which indicates that deconjugation by Cap3 is scarless (Extended Data Fig. 9d). In other words, Cap2-dependent conjugation and Cap3-dependent deconjugation can exist in equilibrium, with the cyclase shuttling between different forms.

### Activity of the *B. cereus* CD-NTase *in vitro*

CBASS cyclases are associated with a variety of effector proteins that bind the nucleotide and induce the cellular response such as degradation of DNA, depletion of NAD and membrane disruption among others ^5,7,39^. We observed earlier that the *Bce* Nuc-SAVED effector (Cap4) was activated by synthetic (3’, 3’, 3’) cA_3_ (Fig. 4d), but note that CD-NTases are capable of making a variety of cyclic nucleotides with both 3’-5’ and 2’-5’ linkages ^5^. To investigate the activity of the cyclase further, we incubated the purified cyclase protein with two different ATP concentrations and analyzed the products by HPLC (Fig. 5c, Extended Data Fig. 10a,b). While the cyclase formed mainly cA_3_ with the low ATP concentration (0.2 mM), no cA_3_ peak appeared when the cyclase was incubated with higher ATP concentration (5 mM). However, a minor peak observed with low ATP became more prominent. To investigate which of these products activated the CBASS effector Nuc-SAVED, we performed a plasmid cleavage assay with the respective samples (Fig. 5d, Extended Data Fig. 10d). Only the cyclase product containing mainly cA_3_ (0.2 mM ATP sample) was able to induce plasmid cleavage through activation of Nuc-SAVED. Nuc-SAVED was not activated by the other products from the 5 mM ATP reaction, showing that cA_3_ is the activator of the *Bce* CBASS effector. Since the 2’-5’ and 3’-5’ linked forms often co-elute on HPLC ^5^, we could not discriminate between the two isomeric forms of cA_3_.

We quantified the levels of production of cA_3_ in the low-ATP reaction samples using a standard curve generated with synthetic cA_3_ (Extended Data Fig. 10c). The wild type cyclase generated cyclic nucleotide at a very low level, with less than a single turnover per hour (0.33 and 0.24 cA_3_ h^-1^ calculated from two time points) (Fig. 5e). This data suggests that the wild type *Bce* cyclase is inactive *in vitro*. To investigate this further, we created a variant of the cyclase lacking residues 302-331 that comprise the flexible tail, which cannot be conjugated, and compared the activity of the truncated protein with the wild type cyclase and with the cyclase conjugated to the PspA protein, purified from *E. coli* (Fig. 5e, Extended Data Fig. 9b). Cyclase activity was measured by assaying the activity of the Nuc-SAVED effector in a fluorescent assay (analogous to ^40^) where the release of a fluorescent signal, generated by cleavage of a 30 bp long double-stranded DNA molecule containing a fluorescent probe and a quencher, was followed. Using this highly sensitive assay, we observed that all three cyclase species: conjugated, unconjugated and truncated, activated the Nuc-SAVED effector to the same extent, within error (Fig. 5f). Thus, our data suggest that the unconjugated, truncated, and PspA-conjugated cyclase species are all inactive forms, suggesting that priming requires deconjugation from PspA by Cap3, and subsequent as yet undetermined steps, to activate the cyclase and initiate the CBASS response.

## Discussion

The regulation of nucleotide cyclases involved in antiviral defence is a key aspect of these innate immunity pathways in both bacteria and eukarya. Recent studies of type II CBASS systems have uncovered the important roles for Cap2-mediated conjugation and Cap3-mediated deconjugation of the cyclase, but left several mechanistic questions unanswered ^22,23,38^. Our investigation of the type II CBASS system from *B. cereus* shows that Cap2 conjugates the cyclase to the major effector of the Psp response, PspA. This posttranslational modification is reversible by Cap3. Thus, to prevent activation of CBASS in the absence of phage, and hence a drastic downstream response, CBASS uses its protein modification machinery to conjugate the cyclase to the highly conserved PspA protein. To the best of our knowledge, this is the first example of a CBASS system using cellular components outside of the defence system to regulate immunity. Regulation of CBASS immunity can be achieved on several levels and the conjugation of cyclase in the absence of phage to a highly conserved and abundant protein like PspA adds a novel aspect to the picture. It has been shown before that the eukaryotic homolog cGAS is activated by DNA-induced oligomerisation ^41–43^. Interestingly, a similar phenomenon has been described very recently for the cyclase enzyme of a short Type II CBASS system. In this case, clustering is achieved by poly-ligation of individual cyclase monomers onto each other ^23^, reminiscent of poly-ubiquitination. We hypothesize that by ligating the cyclase to a highly abundant protein, such clustering could be prevented. Whether conjugation to PspA is solely preventing clustering, and thus activation of the cyclase, or has an additional physiological role can only be speculated at the moment. The ability of Psp systems to respond to phage infection by sensing cell envelope damage could add another level of regulation to CBASS systems, potentially regulating Cap3-mediated deconjugation and, thus, activation of the survival program. The observed association of the *Bce* cyclase to bacterial membranes, a common feature shared with eukarytotic cGAS homologs, gives new insights into the cellular localization of CBASS. Whether membrane association plays a secondary role apart from positioning the cyclase in close proximity to PspA rods, can only be speculated at the moment.

The role of the Cap3 enzyme has been a point of speculation. It clearly can deconjugate CD-NTases from a range of partner proteins ^22,38^. When a range of phages were used to challenge an *E. coli* CBASS system, Cap3 was essential for immunity against phages T4 and T6 but required for phages lambda and T2 ^38^. In contrast, when a *V. cholerae* CBASS system was tested in a heterologous *E. coli* host, Cap3 was not required for immunity against T2, T4, T5 or T6 phages ^22^. In fact, deletion of Cap3 was associated with increased levels of cGAMP production by the *V. cholerae* cyclase immunoprecipitated from *E. coli* cells, leading the authors to propose a model whereby Cap3 is an antagonist of Cap2 that limits cyclase priming ^22^. Finally, Cap3 was not required for antiphage immunity in a *P. aeruginosa* system ^17^. In contrast, the Cap3*^Bce^* enzyme increases immunity provided by the *Bce* CBASS system, as observed by improved defence against phages (Fig. 1b,c).

Our data suggests that the *Bce* Type II CBASS system functions by providing the cell with inactivated, Psp-conjugated cyclase, which is readily available in the event of infection. The observed inhibition of the *Bce* cyclase at ATP levels typical for normally growing bacterial cells could have a regulatory function *in vivo.* In fact, it has been described previously that phage infection affects intracellular nucleotide levels ^44^, a disturbance possibly affecting the activity of the cyclase and/or the Cap2 protein. Whether these observations compose a multi-level regulation of the cyclase, or if they evolved independently, will be interesting to study in the future.It has been observed before that CBASS, in contrast to other defence systems, responds rather late during the infection cycle of the phage ^3^. Our observations support this idea as we detected cA_3_ only 30 min after infection (Extended Data Fig. 6). After the same time, we detected a decrease in the amount of the cyclase-PspA conjugate in infected cells (Fig. 4a,b). Based on this and our observation of earlier accumulation of the second messenger in cells lacking PspA (Fig. 4e), we suppose that deconjugation of the cyclase is necessary for activation.

A working model for the regulation of the *B. cereus* type II CBASS is presented in Figure 6. In uninfected cells, newly synthesised cyclase is conjugated by Cap2 to the abundant cellular protein PspA. An equilibrium between Cap2-mediated conjugation and Cap3-mediated deconjugation may be established, with the majority of the cyclase conjugated. This could be an effective means to prevent cyclase activation by oligomerisation – a mechanism that appears conserved in both eukaryal and bacterial systems ^23,41,43^. At later time points following phage infection, Cap3-mediated deconjugation of cyclase from PspA is observed, implying a role for Cap3 in sensing infection. It is possible that this relates to the changes in PspA organisation apon phage infection ^24,31^, making deconjugation more favourable. The release of cyclase from its conjugation partner could then allow Cap2-mediated activation of the cyclase, possbily due to oligomerisation of multiple cyclase molecules and concomitant activation, as observed for the type I CBASS system ^23^. PspA is near-universal in bacteria ^45^, but was not observed as a conjugation partner in studies conducted in *E. coli* ^22,38^. The use of over-expression platforms and heterologous hosts, while essential for initial biochemical studies, can mask important aspects of these defence systems. Equally, alternative conjugation partners may be utilised in preference over PspA in other bacterial lineages. Finally, we have not directly observed the activated cyclase in the *B. cereus* system, so this remains a key point for future study.

**Figure 6.**
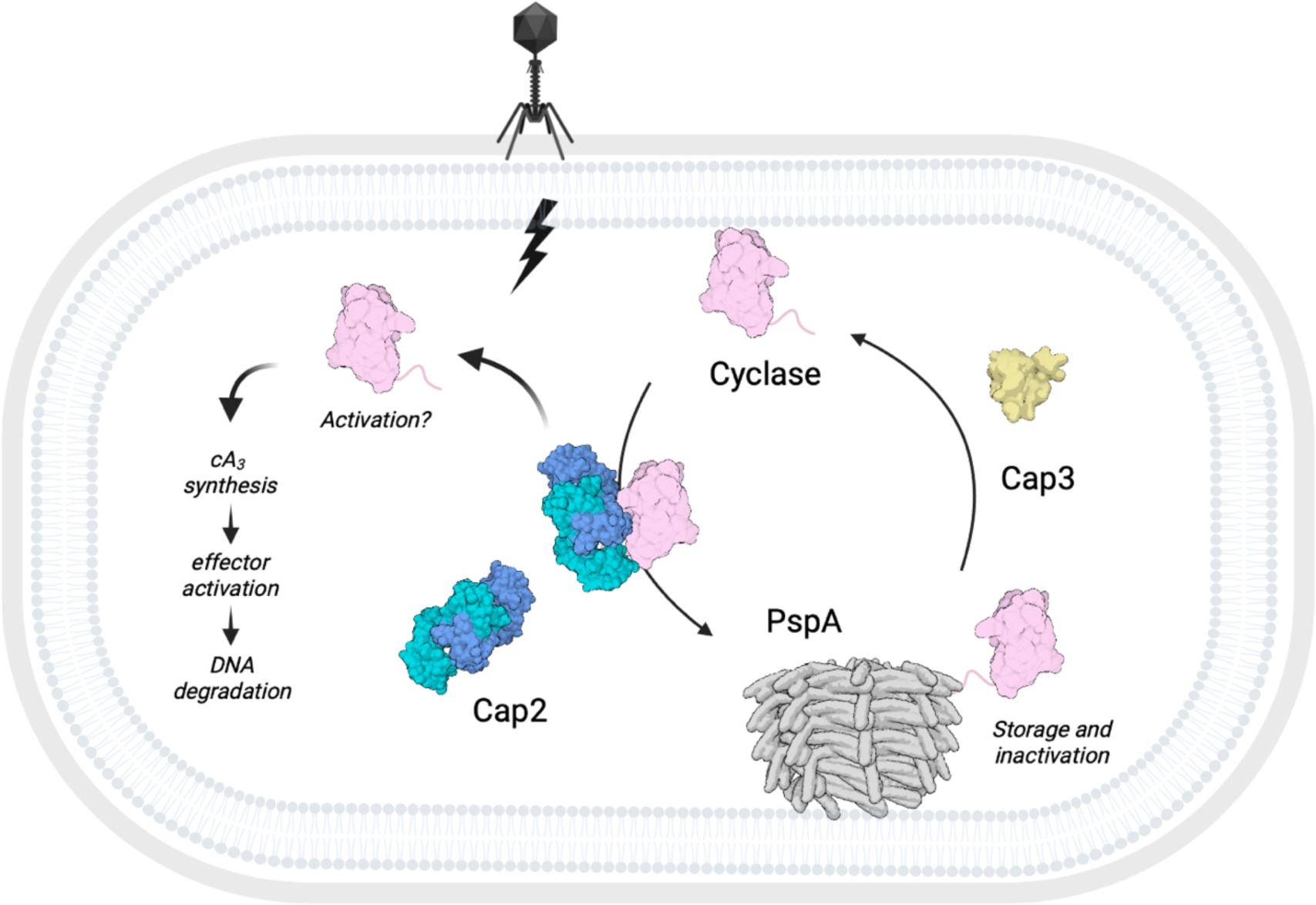
Model of Type II CBASS regulation through cyclase conjugation. In the absence of phage infection, Cap2 ligates the cyclase to the phage shock protein A (PspA) to prevent activation and induction of CBASS. The cyclase-PspA conjugate is cleaved by the isopeptidase Cap3. This equilibrium of unconjugated/conjugated cyclase is shifted towards unconjugated cyclase upon phage infection. Activation of the cyclase is Cap2-dependent and leads to degradation of dsDNA through activation of the CBASS effector Nuc-SAVED. Created with BioRender.com.

## Acknowledgements

This work was funded by a European Research Council Advanced Grant (grant number 101018608) to MFW. LK was funded by an EMBO postdoctoral fellowship (grant number ALTF 234-2022). LG was funded by the UKRI Biotechnology and Biological Sciences Research Council (BBSRC) (grant number BB/T00875x/1). We would like to express our gratitude to Prof. Dr Jörg Stülke for generously providing *Bacillus subtilis* strains, plasmids and antibodies α-HPr and α-Rny. We wish to thank Dr Clarissa Czekster and Dr Gary Tang for granting us access to the HPLC instrument and their advice. We would like to thank Katharina Kohm and Veronika Lutz for the initial isolation of the new phage isolates. Prof. Dr Fabian Commichau is acknowledged for providing training in phage biology.

## Author contributions

LK and MFW conceived and designed the study. LK, LG, and SG performed the experiments. LK, LG and MFW analyzed the data. SS performed the mass spectrometry analysis. RH isolated the phages. MFW acquired funding. LK wrote the original draft of the manuscript. All authors reviewed and edited the manuscript.

## Additional information

Supplementary information is available for this paper. Correspondence and requests for materials should be addressed to Malcolm F White or Larissa Krüger.

## Methods

### Strains, media and growth conditions

*E. coli* DH5α, and BL21 (DE3) or C43 ^46^ were used for cloning and for the expression of recombinant proteins, respectively. All *B. subtilis* strains used in this study are derivatives of the laboratory strain 168. *B. subtilis* and *E. coli* were grown in Lysogeny Broth (LB) or in sporulation (SP) medium ^46,47^. For growth assays and the *in vivo* interaction experiments, *B. subtilis* was cultivated in LB. The media were supplemented with ampicillin (100 µg/ml for *E. coli*), kanamycin (50 µg/ml for *E. coli*, 10 µg/ml for *B. subtilis*), chloramphenicol (35 µg/ml for *E. coli*, 5 µg/ml for *B. subtilis*), erythromycin and lincomycin (2 and 25 µg/ml, respectively), or Zeocin™ (35 µg/ml) if required.

### Isolation of novel *Bacillus subtilis* phages

The laboratory strain *Bacillus subtilis* Δ6 ^27^, a derivate of *B. subtilis* 168, was used as host for the isolation of novel phages as described previously ^48^. Shortly, raw sewage from a municipal sewage treatment plant in Göttingen, Germany served as environmental phage source and was cleared by centrifugation at 5000 g for 10 min, following filtration through a sterile filter (0.45 μm; Sarstedt). The host was infected by mixing 100 μl of an overnight culture with 2 ml of processed sewage water, followed by 5 min incubation at room temperature to allow adsorption. This bacterial suspension was then mixed with 2.5 ml of 0.4 % agarose (50 °C) (Fisher BioReagents™) dissolved in LB medium, spread on a LB plate, and incubated overnight at 28 °C. Individual plaques were picked, serially diluted and higher dilutions were used for re-infection of the host. Cells were grown until complete lysis and the supernatant processed as described above. This re-infection was repeated to scale up the phage lysate. For genome sequencing of the novel isolates, phage DNA was isolated as described before ^49^. Sequencing was performed using an Illumina MiSeq sequencer (Illumina). Phages were classified according to nucleotide sequence homology to the closest relative as revealed by Megablast within the Geneious Prime® 2022.2.1 software. The genomic sequences of the new phages and their closest relative (downloaded from NCBI on 10 March 2023) were aligned using MMseqs within the pyGenomeViz package (https://github.com/moshi4/pyGenomeViz, accessed on 10 April 2023; input: gbk files; mode: pgv-mmseqs). A list of all phage isolated in this study can be found in Supplementary Table 1.

### Phage amplification and storage

Phage lysates were generated from *B. subtilis* Δ6 using the modified double agar overlay plate technique. For this, a culture of *B. subtilis* Δ6 was precultured to late exponential growth phase in LB medium. 100 µl of bacterial cells were mixed with 100 µl phage dilution, incubated for 5 min at RT to allow adsorption of the phage, and mixed with 2.5 ml LB supplemented with 0.4 % agarose (50 °C), and poured over a prewarmed (37 °C) LB plate. Plates were incubated overnight at 37 °C. Phages were harvested on the following day by adding 5 ml of LB medium directly onto the plate, incubating for at least 30 min at room temperature, and collecting and filtering the resulting liquid through a 0.2 µm Nanosep filter. Lysate titers were determined by infection of *B. subtilis* Δ6 using the modified double agar overlay plate technique as described above and the resulting PFU/ml was calculated. Phage lysates were stored at 4 °C.

### Phage infection assays

Phage infection assays were performed on plates and in liquid culture. The desired *B. subtilis* strains were precultured over night at 28 °C and used to inoculate fresh LB medium. This culture was grown until late exponential growth phase. For infection on plates, 100 µl were mixed with 2.5 ml LB supplemented with 0.4 % agarose (50 °C) and poured over a prewarmed LB plate. Serial dilutions of the phage lysate were spotted onto this plate. Plates were incubated overnight at 37 °C and pictures were taken on the following day. For quantification, PFU/ml was calculated from at least three biological replicates. Fold defence was calculated as the ratio between the PFU/ml obtained from wild type (no CBASS) and strains containing the defence system. For infection in liquid medium, the optical density of the culture at 600 nm (OD_600_) was adjusted to 1.0 and the cells were used to inoculate a 96 well plate (Microtest Plate 96 Well, Sarstedt) containing LB medium. Growth was tracked in an Epoch 2 Microplate Spectrophotometer (BioTek Instruments) at 37 °C with linear shaking at 237 cpm (4 mm) for 20 h, and an OD_600_ was measured in 2 min intervals.

### DNA manipulation

Transformation of *E. coli* and plasmid DNA extraction were performed using standard procedures ^46^. All commercially available plasmids, restriction enzymes, T4 DNA ligase and DNA polymerases were used as recommended by the manufacturers. Chromosomal DNA of *B. subtilis* was isolated as described ^47^. *B. subtilis* was transformed with plasmid and genomic DNA according to the two-step protocol ^47^.

### Construction of mutant strains by allelic replacement

Deletion of the *pspA* gene was achieved by transformation of *B. subtilis* Δ6 with a PCR product constructed using oligonucleotides to amplify DNA fragments flanking the target genes and an appropriate intervening resistance cassette as described previously ^50^. The integrity of the regions flanking the integrated resistance cassette was verified by sequencing PCR products of about 1,100 bp amplified from chromosomal DNA of the resulting mutant strains. A list of all strains constructed in this study can be found in Supplementary Table 2.

### Plasmid constructions and mutagenesis

The wild type CBASS operon from *B. cereus* WPysSW2 was ordered as genomic DNA from IDT and the desired genes were amplified using appropriate oligonucleotides that attached specific restriction sites to the fragment. Plasmids derived from pGP1460 (integration into *lacA*) ^51^ were linearized with ScaI for genomic integration. The integrity of the integration was confirmed by PCR and subsequent sequencing of the region.

For protein expression in *E. coli*, the genes encoding CD-NTase, Cap2, Cap3 and Nuc-SAVED were codon optimized and obtained from Integrated DNA Technologies (IDT). Each gene was subcloned into the vector pEhisV5TEV ^52^, allowing expression with a cleavable N-terminal polyhistidine tag, or into pGP172 for expression with an N-terminal Strep-tag. The *pspA* gene was amplified from *B. subtilis* 168 chromosomal DNA using oligonucleotides that attached specific restriction sites to the fragment and cloned into the vector pEhisV5TEV. The PspA mutants K147 and K220 were generated by site directed mutagenesis of the *pspA*-pEhisV5TEV vector. For co-expressions in *E. coli*, the CD-NTase remained within the pEhisV5TEV vector to retain cleavable polyhistidine tag, whilst other proteins (Cap2, Cap3 and PspA) were expressed on pCDFduet and pACYCduet. All synthetic genes, plasmids and primers are listed in Supplementary Table 3, 4 and 5.

### Protein expression and purification

*E. coli* C43 (DE3) was transformed with the plasmid pEhisV5TEV encoding the wild type *Bce* cyclase or a variant truncated after amino acid 301, Nuc-SAVED, Cap2, Cap3, wild type PspA, or the mutants, or pGP172 encoding Strep-Cap2. Expression of the recombinant proteins was induced by the addition of isopropyl 1-thio-β-D-galactopyranoside (final concentration, 0.4 mM for pEhisV5TEV or 1 mM for pGP172) to exponentially growing cultures (OD_600_ of 0.8) of *E. coli* carrying the relevant plasmid. Cell pellets were resuspended in lysis buffer (50 mM Tris-HCl, 250 mM NaCl, 10 mM Imidazole, 10 % glycerol, pH 7.5) supplemented with 1 mg/ml lysozyme and protease inhibitor (Roche). The cells were lysed by sonication on ice (MSE Soniprep 150). After lysis, the crude extract was centrifuged at 117,734 x g for 30 min and then passed over a Ni^2+^nitrilotriacetic acid column (His-trap FF crude, Cytiva). Ni^2+^nitrilotriacetic acid columns were washed with wash buffer (50 mM Tris-HCl, 250 mM NaCl, 30 mM Imidazole, 10 % glycerol, pH 7.5) to clear unbound protein. Target protein was eluted with an imidazole gradient or biotin (5 mM). After elution, the fractions were tested for the desired protein using SDS-PAGE. To remove the 8x-His-TEV tag from the proteins, the relevant fractions were combined, and the tag was removed with the TEV protease (ratio 10:1 w/w) during overnight dialysis (Biodesign™ Cellulose Dialysis Tubing Roll, 10 kDa) against wash buffer. The cleaved TEV moiety and the protease were removed using a fresh Ni^2+^nitrilotriacetic acid column. The purified protein was concentrated in a Merck Amicon™ Ultra-15 Centrifugal Filter device (cut-off 10 kDa). The protein was loaded on a HiLoad 16/600 Superdex 200 pg column pre-equilibrated with gel filtration buffer (20 mM Tris-HCl, 250 mM NaCl, 10 % glycerol, pH 7.5) and the fractions containing pure protein were collected and concentrated again. The protein samples were stored at −70°C until further use.

Expression of wild type PspA or the mutant proteins was induced with 0.4 mM isopropyl-β-D-1-thiogalactoside (IPTG) and cells grown for 4 h at 25 °C. PspA was purified as described above with the exception that the Ni^2+^nitrilotriacetic acid column was washed with 20 column volumes (CV) of wash buffer containing 50 mM Tris-HCl pH 7.5, 500 mM NaCl, 30 mM imidazole, 10 % glycerol, then 4 CV of 50 mM Tris-HCl pH 7.5, 500 mM NaCl, 50 mM imidazole, 10 % glycerol and then eluted directly with 6 CV of elution buffer containing 50 mM Tris-HCl pH 7.5, 500 mM NaCl, 500 mM imidazole and 10 % glycerol. The eluted peak was pooled and concentrated using an Amicon Ultra-15 (Millipore) centrifugal filter (MW cut-off 10 kDa) prior to loading onto an equilibrated HiTrap desalting column (Cytiva), then washed with 50 mM Tris-HCl pH 7.5, 500 mM NaCl, 30 mM imidazole, 10 % glycerol. The desalted protein was collected and treated with TEV protease for 2 h at room temperature, with gentle agitation. The protein was isolated from TEV protease by passing through another HisTrap FF crude column and the unbound fraction collected and concentrated. PspA was further purified by size exclusion chromatography (S200 16/60, Cytiva) in buffer containing 20 mM Tris pH 7.5, 500 mM NaCl, 10 % glycerol. Chosen fractions were concentrated and aliquoted, then frozen at −70 °C.

Strep-tagged proteins were expressed, lysed and ultracentrifuged as His-tagged proteins above, in buffer W (100 mM Tris-HCl, 150 mM NaCl, pH 8.0), unless stated otherwise. The crude extract was then applied to StrepTactin column (IBA, Göttingen, Germany) and the column washed with 100 mM Tris-HCl, 150 mM NaCl pH 8.0. The protein was eluted with a biotin gradient (5 mM). Relevant fractions were identified by SDS-PAGE and concentrated using Vivaspin turbo 15 (Sartorius) prior to storage as described above.

### Co-expression and purification of the cyclase*^Bce^-*PspA*^Bsu^* conjugate

Co-expression strains (His-tagged cyclase*^Bce^*, untagged PspA*^Bsu^* and Cap2*^Bce^*) were generated through transformation of *E. coli* C43 (DE3) with pV5SpHISTEV, containing wild type cyclase*^Bce^*, or the truncated variant (1-301), in addition to the dual-expression vector pCDFduet that contained PspA*^Bsu^* and Cap2*^Bce^*. Cultures were induced for protein expression with 0.4 mM IPTG for 4 h at 25 °C, conditions previously identified as optimum for cyclase-PspA conjugate formation ^52^. Cyclase-PspA conjugate was purified following His-tagged protein method detailed above. As the only protein with His-tag present was the cyclase, minimal Cap2/PspA were expected unless conjugated to/interacting with cyclase. Multiple cyclase-PspA conjugation products were separated through SEC on HiLoad 26/600 Superdex 200 pg column, flow rate of 1.5 ml/min for increased separation of varying MW species. Fractions containing individual conjugation species were then identified via SDS-PAGE and pooled separately prior to concentrating and storing in −70 °C as above.

### Pulldown of cyclase*^Bce^* in *E. coli*

Co-expression strains of the His-cyclase and PspA with or without additional expression of Cap2 or Cap3 were grown in 10 ml LB and induced during exponential growth phase with 0.4 mM IPTG. Expression strains were then incubated for 4 h 25 °C and cells harvested by centrifugation. His-tagged cyclase was purified and visualised on SDS-PAGE as described previously ^52^. Briefly, cell pellets were resuspended in lysis buffer (50 mM Tris-HCl, 250 mM NaCl, 10 mM Imidazole, 10 % glycerol, pH 7.5), before being lysed by sonication on ice. The cell lysate was then centrifuged at 10,000 x g for 10 min before supernatant was added to Biosprint plasticware. His-tagged proteins were isolated using Nickel beads (His Mag Sepharose Ni, GE Healthcare) in the QIAGEN BioSprint machine, with wash and elution buffers containing 50 mM Tris-HCl, 500 mM NaCl, 10 % glycerol, pH 7.5, and 30/ 500 mM imidazole, respectively. Eluted His-tagged protein and total protein containing fractions were analysed by SDS-PAGE and Western blot.

### Purification of cyclase*^Bce^* from *B. subtilis* and separation of cell extract

For overexpression of the cyclase in *B. subtilis*, strain LK40 was transformed with either pLK20 (Strep-cyclase), pLK21 (Strep-cyclase^ΔC^), or the empty vector control (pGP380). For comparison of levels of cyclase conjugate the *B. subtilis* strains LK06, LK10, LK18, LK20 and LK51 which contained single genomic integrations of the CBASS operon were used. The respective strains were cultivated in LB medium until exponential growth phase was reached (OD_600_ ∼ 0.5). If desired, cells were infected with the phage Goe23 and incubated for another 5, 15, or 30 min. The cells were harvested by centrifugation at 5,000 rpm and 4 °C and the pellets stored at −20 °C. For separation of cytosolic and membrane fractions of the crude extract, the pellets were resuspended in buffer M (50 mM NaH_2_PO_4_, 50 mM Na_2_HPO_4_, pH 6.8) supplemented with 1 mg/ml lysozyme and protease inhibitor (Roche) and opened by sonication. To remove cell debris and unbroken cells, the lysate was centrifuged at 5,000 rpm for 20 min at 4 °C. The supernatant was subjected to ultracentrifugation at 21,000 g for 90 min at 4°C to pellet the membranes. The supernatant, containing cytosolic proteins, was removed, the pellet washed with buffer M and the membranes pelleted again in an additional centrifugation step (21,000 g, 60 min, 4 °C). The supernatant was discarded and the wash step repeated one more time. The resulting pellet, containing the membranes, was resuspended in buffer M. To achieve solubilization of the membranes, the fraction was 1:1 diluted with buffer M containing 0.5 % DDM (final concentration 0.25 %) and incubated rotating over night at 4 °C. Protein concentration of cytosolic and membrane fractions was determined and equal amounts of the fractions were separated on an SDS gel and analyzed by staining with Coomassie and Western blot analysis. To confirm the proper separation of cytosolic and membrane fractions, membranes were also developed with α-HPr and α-Rny primary antibodies as cytosolic and membrane protein control, respectively. The cyclase was further purified from both cytosolic and membrane fractions using Protein A coupled Dynabeads (Invitrogen™) that were saturated with α-cyclase antibody for immunoprecipitation. The eluates were separated by SDS-PAGE and analyzed by Western blot. Bands of interest were subjected to further analysis by mass spectrometry.

### Western blot analysis

Polyclonal antibodies against the *B. cereus* cyclase or the PspA protein from *B. subtilis* were produced in rabbits (Kaneka Eurogentec S.A) and the anti-cyclase antibody was purified with Cyanogen bromide-activated-Sepharose® (Merck) by immobilizing purified cyclase. The purified α-cyclase antibody was used at 1:10,000 dilution for detection and the α-PspA serum at 1:5,000 dilution. Western blot analysis was performed on cell extracts and purified protein to follow the modification of the cyclase protein. Samples were resolved using SDS-PAGE and transferred to PVDF membranes. Membranes were blocked for at least 1 h at RT in blocking buffer (1x PBS, 5 % w/v milk powder, 0.025 % Tween20). Incubation with the respective antibodies (α-cyclase, α-HPr, α-Rny) happened over night at 4 °C. Blots were extensively washed and incubated with the appropriate α-Rabbit secondary antibodies (Licor infrared (800CW/680RD)) at 1:20,000 dilution in blocking buffer. The specific protein bands were visualized using the Licor Odyssey CLx and images adjusted using the software ImageJ (http://rsb.info.nih.gov/ij/index.html). Bands were analysed and quantified with ImageJ by comparing the intensity of selected bands to a standard band on the same blot.

### Conjugation assay

For reconstruction of the conjugation *in vitro*, the cyclase (5 µM) was incubated together with Cap2 (5 µM), PspA (10 µM), 10 mM MgCl_2_ and 0.3 mM ATP in 100 mM Tris-HCl, 150 mM NaCl, pH 8.0 for 30 min at 25 °C. The samples were separated by SDS-PAGE and analysed by Coomassie staining or Western blot with an antibody raised against the cyclase.

### Mass spectrometry analysis

Protein bands were excised from the gel and prepared for mass spectrometry analysis using established protocols ^53^. Briefly, this included destaining with ethanol:water, reduction with dithiorethritol and alkylation with iodoacetamide. Samples were digested with either trypsin, AspN, or a combination of trypsin and GluC. The peptides were soaked from the gel pieces, and the eluent concentrated to 20 µl. The samples were analysed by nanoLCMSMS on a ThermoScientific Fusion Lumos Orbitrap mass spectrometer coupled with a ThermoScientific u3000 nanoLC. A LC was configured in trap elute format, with the Acclaim Pepmap 100 100 µm x 2 cm nanoViper trap and the Pepmap RSLC C18 3 µm 100A 75 µm x 15 cm easyspray column, both ThermoScientific. 5-10 µl of sample was injected onto the trap in 15 µl/min of loading buffer (0.05 % Trifluoroacetic acid in water), and run for three minutes. The trap was switched in line with the analytical column and the sample eluted at a gradient over 65 min (A = 100 % water with 0.1 % formic acid, B = 20 % water 80 % acetonitrile, 0.1 % formic acid, 2 % A to 3 min, linear to 40 % A over 42 min, linear to 95 % A over 4 min, hold for 5 min, linear back to 2 % A, and re-equilibrate for 10 min). The flow from the column was sprayed directly into the Easyspray orifice at a voltage of 1700 V positive ionisation. Mass spectrometry data was collected from 350 to 2000 on the orbitrap at resolution 120000 for the survey scan and a cycle time of 2 sec for Data Dependant Acquisition conditions for MSMS on the orbitrap trap at a resolution of 30,000. Both Electron Transfer Dissociation (ETD) and Higher Energy Collison Dissociation (HCD) fragmentation techniques were used. Raw data was exported and extracted using ms convert (ProteoWizard). The data searched using Mascot search engine (MatrixScience) against an internally generated database of 6700 protein sequences containing the sequences of our recombinant proteins, or the *B. subtilis* specific protein database (UniProt Proteome ID UP000001570). Settings were 20 ppm on the MS and 0.1 Da on the MSMS data, with a fixed modification of carbamidomethyl on cysteine and variable oxidation of methionine. For the identification of the conjugation site on PspA, the mass of the last C-terminal tryptic peptide of the cyclase (KPGGFA) minus one water molecule (C(27) H(39) N(7) O(6) 557.296182) was set as a variable modification on lysine residues.

Intact mass measurement was carried out on a Waters Xevo G2S TOF mass spectrometer with Waters Acquity LC. 10 µl of 1 µM sample was desalted on-line through a waters MassPrep On-Line Desalting Cartridge 2.1 x 10 mm, eluting at 200 µl/min, with an increasing acetonitrile concentration (2 % acetonitrile, 98 % aqueous 1 % formic acid to 98 % acetonitrile, 2 % aqueous 1 % formic acid) and eluted directly into the MS, with a lock mass of LeuEnk to ensure stable calibration. The spectra across the elution peak were combined and the charged ion envelope deconvoluted with MaxEnt algorithm using the peak width at half height of the most intense peak, to a resolution of 0.1 Da.

### Cyclase product quantification by HPLC

To obtain cyclase products, the reaction was set up as follows: 50 µM cyclase, 0.2 mM/5 mM ATP in cyclase buffer (50 mM HEPES, 150 mM KCl, 10 mM MgCl_2_, 10 % glycerol, pH 7.5) for 20 min/ 45 min. Reaction samples were filtered through a Pall Nanosep spin filter (3-kDa cut-off) by centrifugation at 13,000 rpm for 10 min to remove protein. 5 µl of product or synthetic standard was injected onto a C18 column (Kinetex EVO P 2.1 × 50 mm, the particle size of 2.6 µm) attached to a Thermo UltiMate 3000 chromatography system. Absorbance was monitored at 260 nm and 40 °C. Gradient elution was performed with solvent A (100 mM ammonium acetate) and solvent B (100 % methanol plus 0.1 % TFA) with a flow rate of 0.3 ml/min as follows: 0-0.5 min, 0 % B; 0.5-3.5 min, 20 % B; 3.5-5 min, 50 % B; 5-10 min, 100 % B. The area under the peak (mAU*min) was determined with the Chromeleon 6.80 software (Dionex). The activity of the cyclase was calculated by fitting the area under the peak values to those of a cA_3_ standard curve.

### Cyclase product quantification by fluorescent Nuc-SAVED assay

Wild type cyclase*^Bce^*, the truncated variant (1-301) or the cyclase*^Bce^*-PspA*^Bsu^* conjugate were incubated with ATP under standardised conditions for comparison of product formation. 20 µM of cyclase was incubated with 250 µM ATP in cyclase buffer (50 mM HEPES, 150 mM KCl, 10 mM MgCl_2_, 10 % glycerol, pH 7.5) for 1 h, unless stated otherwise. Reactions were quenched with 10 mM EDTA, and cyclase denatured 95 °C for 5 min. For rapid quantification of the relative cA_3_ produced by the cyclase variants, a dilution of the cyclase reaction was incubated with the CBASS effector Nuc-SAVED. Method and conditions were adapted from an analogous previously described NucC assay ^40^. The assay contained 50 mM Tris-HCl, pH 7.5, 20 mM NaCl, 10 mM MgCl_2_, 10 %(v/v) glycerol, 100 nM FAM:Iowa Black®double-stranded DNA substrate, 100 nM Nuc-SAVED and 500x dilution of denatured cyclase reaction or 25 nM synthetic cA_3_, unless otherwise stated. The absorbance was measured with a FluoStar Omega plate reader (BMG Labtech) using fluorescence detection (ex/em 485/520 nm) in black, non-binding half-area 96-well plates (Corning). Fluorescence was measured in 30 s intervals at 37 °C. The reaction was initially incubated in the absence of cA_3_/cyclase product for 10 min to measure a base line. The measurement was stopped, synthetic cA_3_/cyclase product added, and the measurement continued for 50 min. A standard curve was generated for Nuc-SAVED with varying synthetic 3’5’ cA_3_ concentrations (0.1 nM – 1 µM) and initial rate of reaction used to generate a standard curve of rate vs [cA_3_]. For the comparison of the different cyclase variants, the cyclase reaction sample was analyzed by SDS-PAGE additionally to normalize the protein amount to the initial rate, as the conjugate sample was less pure compared to the wild type and the truncated cyclase. Bands were analysed and quantified with ImageJ by comparing the intensities of selected bands on the same gel.

### Extraction of nucleotides

For the extraction of nucleotides, *B. subtilis* cultures carrying genomic integrations of the respective CBASS version were grown until exponential growth phase at 37 °C. If indicated, the cultures were infected with phage Goe23 to an MOI of 2 and grown for additional 30 min. A volume of 50 ml of the cultures was harvested immediately, and the pellets resuspended in extraction solution (methanol/water/acetonitrile, 2/2/1). The cells were lysed, and the proteins denatured by vortexing for 20 s and incubation at 95 °C for 10 min. The samples were stored at −70 °C overnight. After thawing, the denatured proteins were separated by centrifugation at 21,000 g and the protein pellet used for determination of the protein amount in each sample. The supernatants containing the cell-free extracts were dried at 40 °C in a Speed-Vac®. The dried pellets were resuspended in 60 µl water and filtered through a Pall Nanosep spin filter (30-kDa cut-off) by centrifugation at 13,000 rpm for 10 min. The filtrate was tested for the CBASS specific nucleotide in a plasmid cleavage assay.

### Plasmid cleavage assay

The assay contained 20 mM HEPES, pH 7.5, 10 mM NaCl, 10 mM MgCl_2_, 10 %(v/v) glycerol, 75 nM plasmid DNA, 1 µM CBASS effector protein Nuc-SAVED and 1.5 µl of the extracted nucleotides or the diluted cyclase reaction product (1:1,000/1:10,000) per 15 µl reaction sample or synthetic cA_3_, unless stated otherwise. The reaction was started by the addition of the nucleotide extracts or the synthetic standard, and the samples incubated at 37 °C for 30 min, for 10 min in case of the cyclase reaction products or in case of the time course experiment for the stated amount of time. The samples were mixed with DNA loading dye (TriTrack, Thermo Scientific) and separated on a 1 % agarose gel (run in 1x TBE). The DNA was stained with SYBR-Green (1;50,000, in 1x TBE) and the gels imaged on Typhoon™ FLA 7000 (Cytiva Life Sciences). The band of the undigested supercoiled plasmid was quantified with ImageJ by comparing the intensity of selected bands to a standard band on the same gel. If indicated, relative values were normalized to the amount of protein extracted in parallel to the nucleotides from the same cultures.

---

**Extended Data Figure 1.**
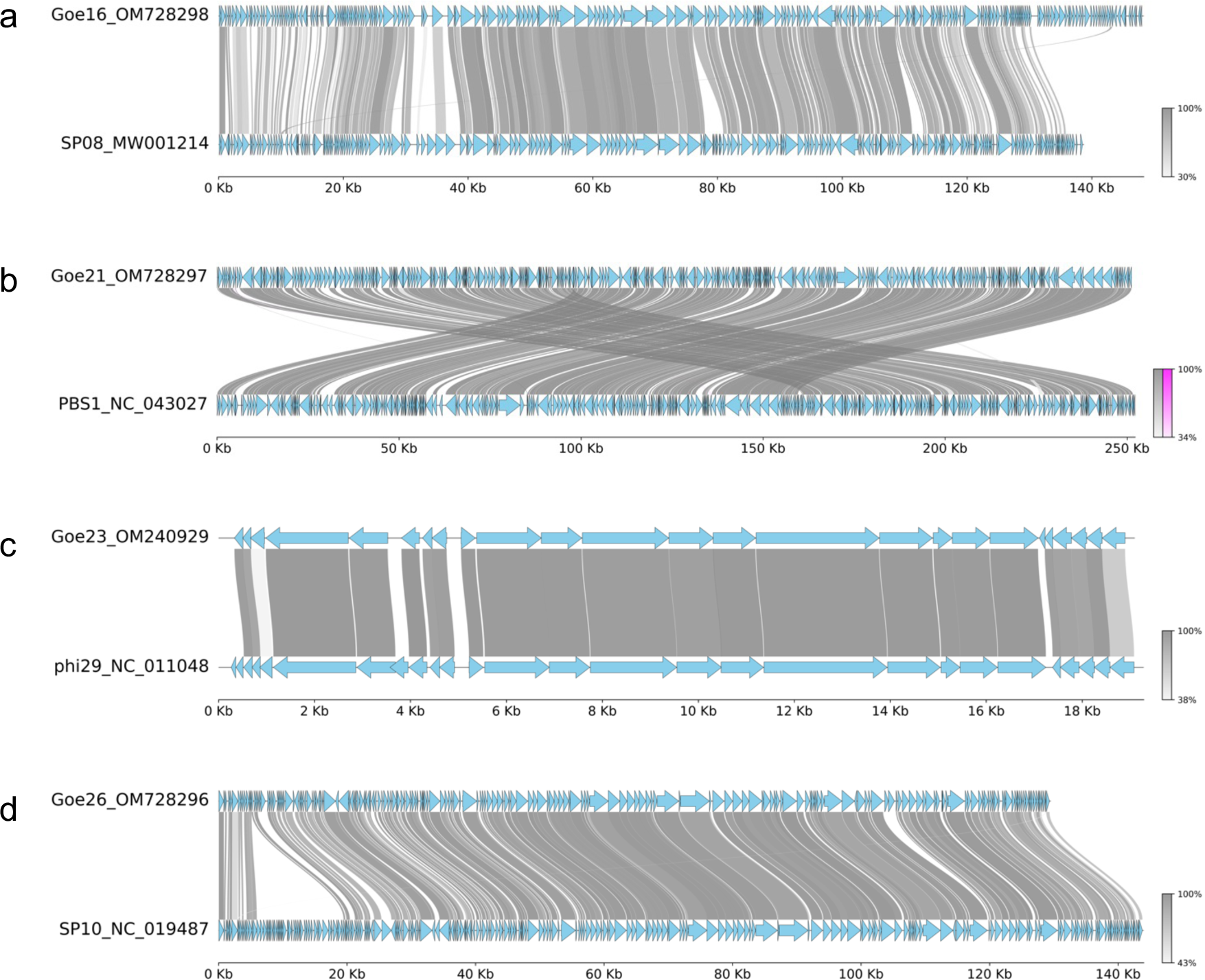
Genome alignments of four new *Bacillus subtilis* phages with their closest relative. The genomic sequences of the novel *Bacillus subtilis* phages Goe16 (a), Goe21 (b), Goe23 (c) and Goe26 (d) were compared to the genome sequence of their closest relative as revealed by Megablast. Genome sequences were aligned with MMseqs within the pyGenomeViz package. blue: CDS; grey: normal link; magenta: inverted link. Color shades represent sequence identities (%).

**Extended Data Figure 2.**
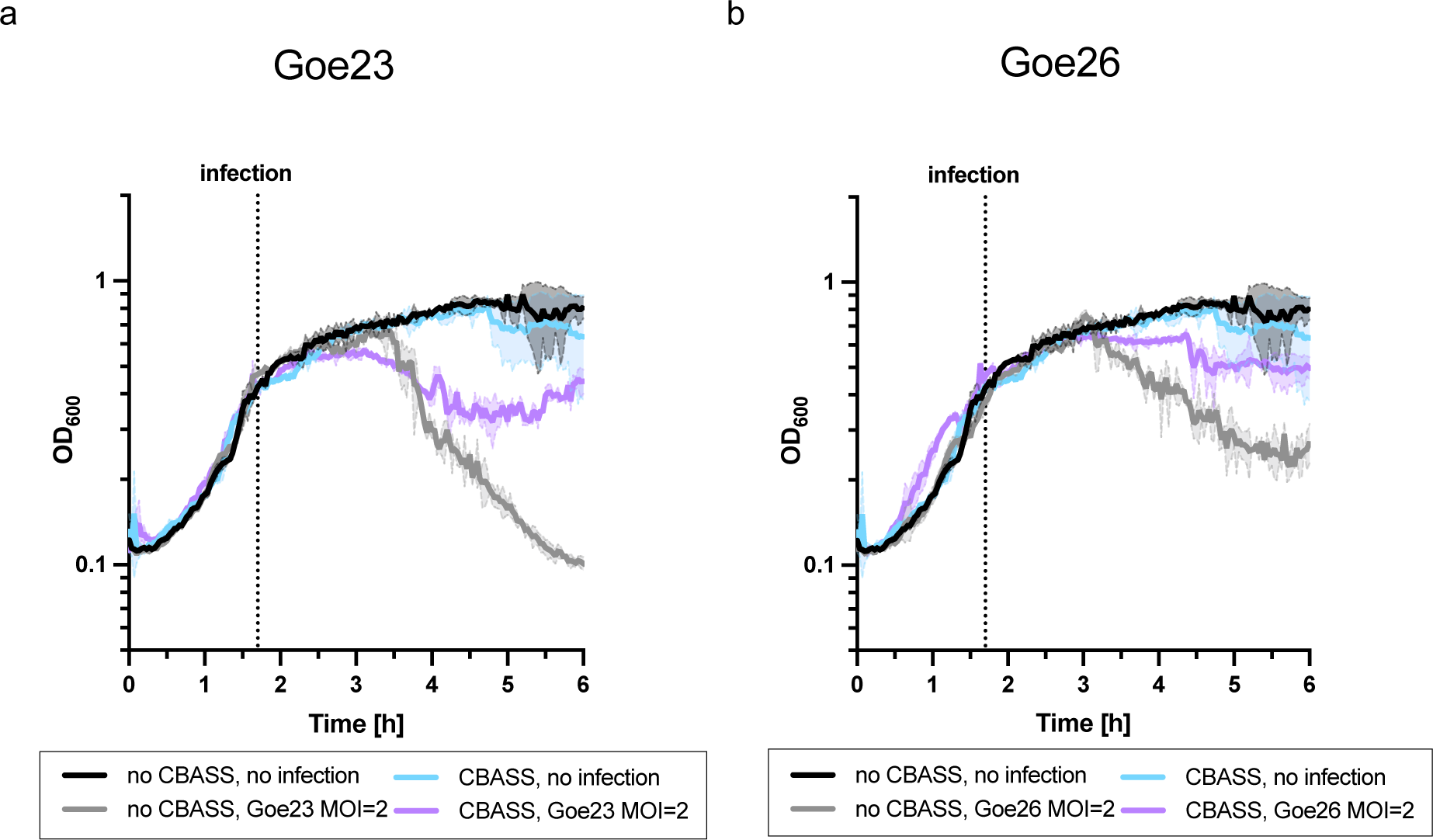
CBASS protects against Goe23 in liquid medium. The CBASS operon of *Bacillus cereus* WPySW2 was integrated into *Bacillus subtilis* Δ6. The growth of wildtype (no CBASS) and cells expressing CBASS was monitored until they reached mid-exponential growth phase and they were infected with phage Goe23 (**a**) or Goe26 (**b**) to an MOI of 2. The experiment was conducted with at least two biologically independent replicates and a representative growth curve is shown.

**Extended Data Figure 3.**
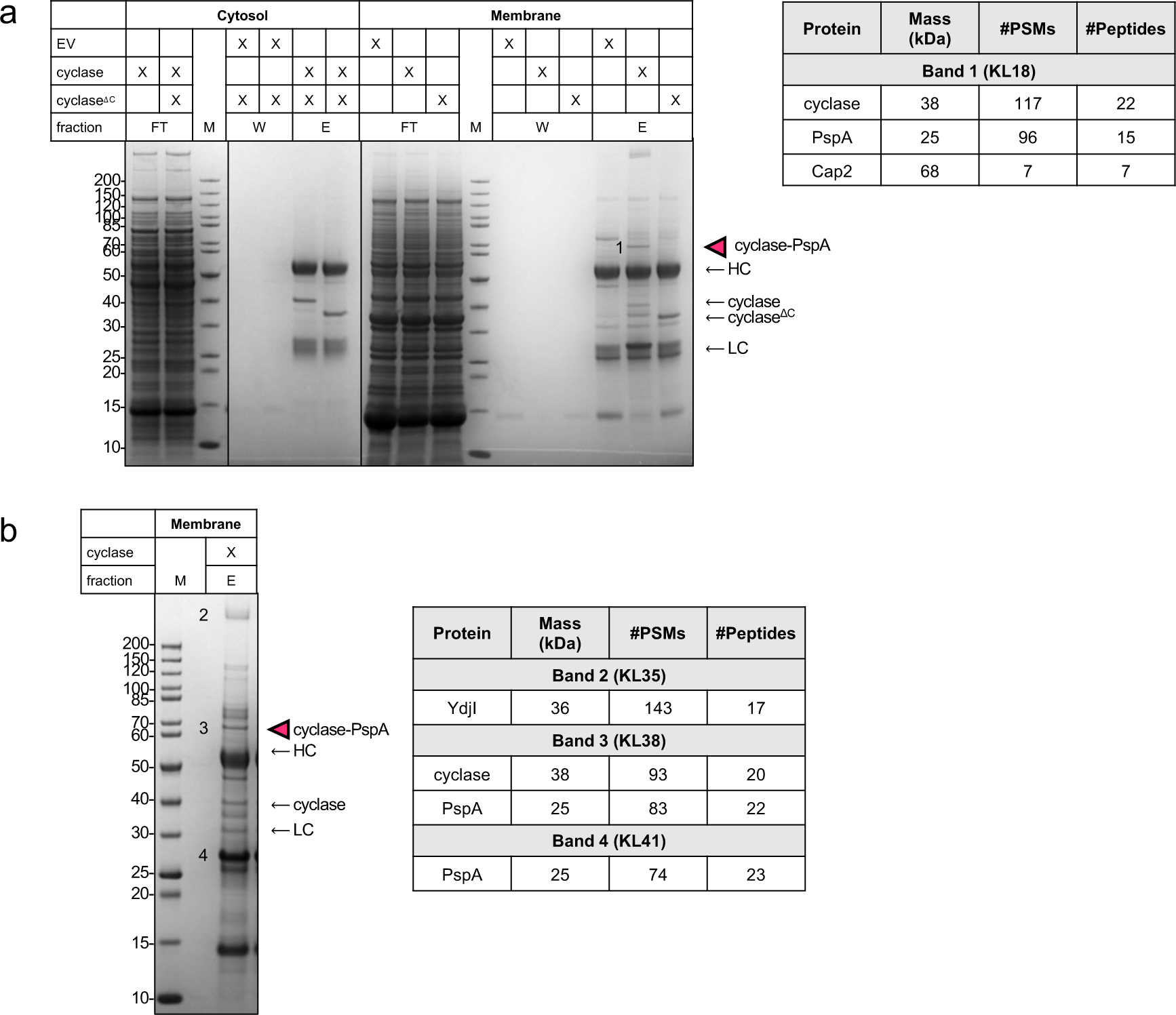
The cyclase is conjugates to the phage shock protein A *in vivo*. **a**, Purification of the overexpressed cyclase (*Bce*) from *B. subtilis* cytosolic and membrane fractions using protein A coupled Dynabeads and a cyclase specific antibody. Abbreviations: EV: empty vector; FT: flow-through; W: wash fraction; E: elution fraction; HC: heavy chain; LC: light chain. Band 1 was excised from the SDS gel, digested with trypsin and analyzed by mass spectrometry. **b**, Replicate of a. Bands were analysed as before. The mass of the last C-terminal tryptic peptide of the cyclase (KPGGFA) was set as a variable modification in the peptide search. The amino acid sequence of PspA is shown and peptides containing the additional mass are emaild and modified lysine residues highlighted in magenta. Abbreviations: PSMs: peptide sequence matches.

**Extended Data Figure 4.**
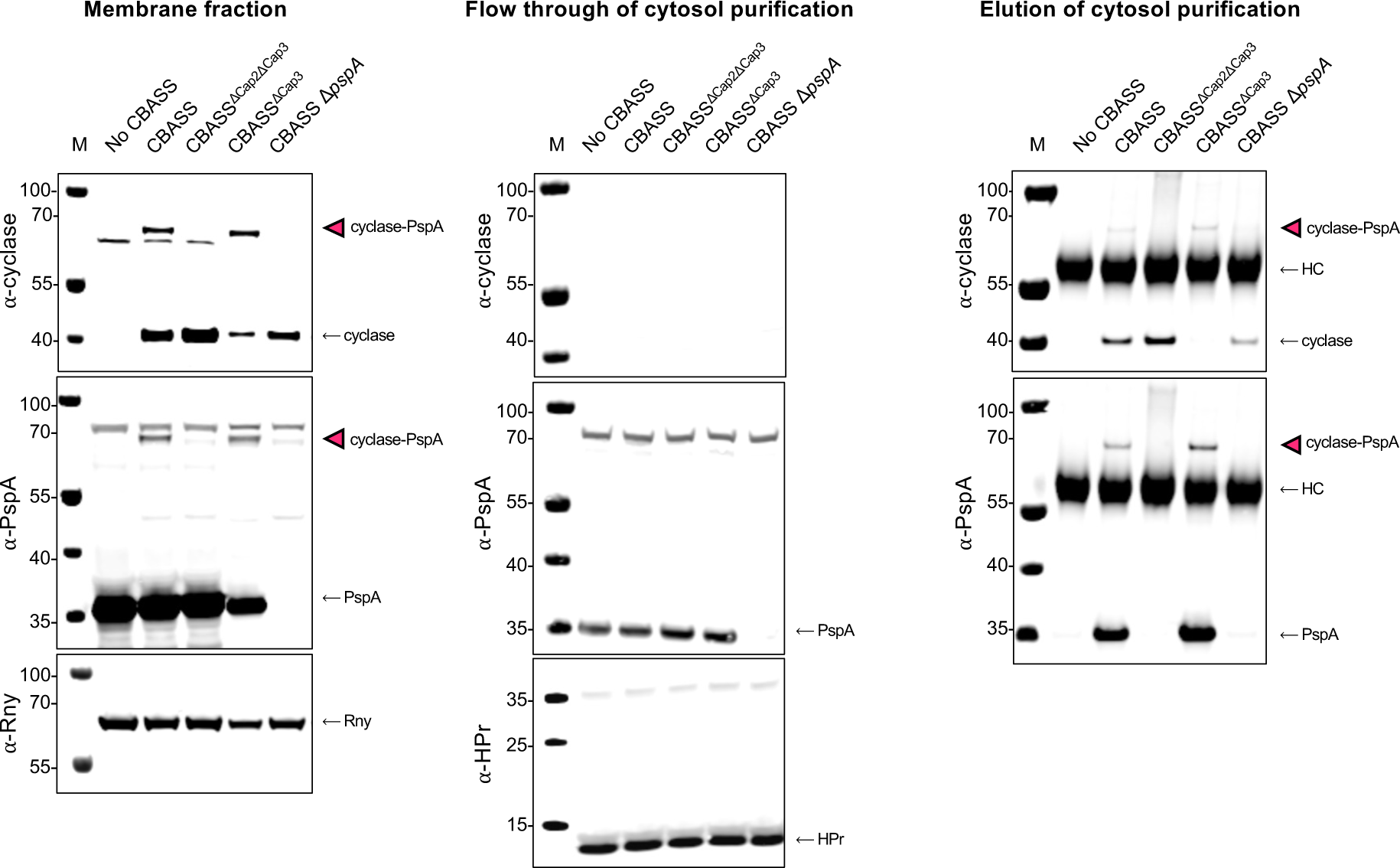
Cyclase conjugation is Cap2 dependent, reversible by Cap3, and PspA is the sole target. *B. subtilis* wild type (no CBASS) and strains expressing genomic integrations of the *B. cereus* CBASS operon were grown and harvested at early exponential growth phase. Cell lysates were separated into cytosol and membranes. Cytosolic fractions were concentrated by purifying the cyclase using protein A coupled beads and a cyclase specific antibody. Membrane fractions, the flow through fractions of the cytosol concentration (loading controls for purification), and the elution fractions were separated by SDS-PAGE and analysed by Western blot with a cyclase- or PspA-specific antibody. Antibodies raised against the RNase Y protein and the phospho-carrier protein HPr served as a loading control for the membrane fraction and cytosol purification, respectively. Abbreviations: HC: heavy chain. The experiment was conducted with two biological replicates and representative images from one replicate are shown.

**Extended Data Figure 5.**
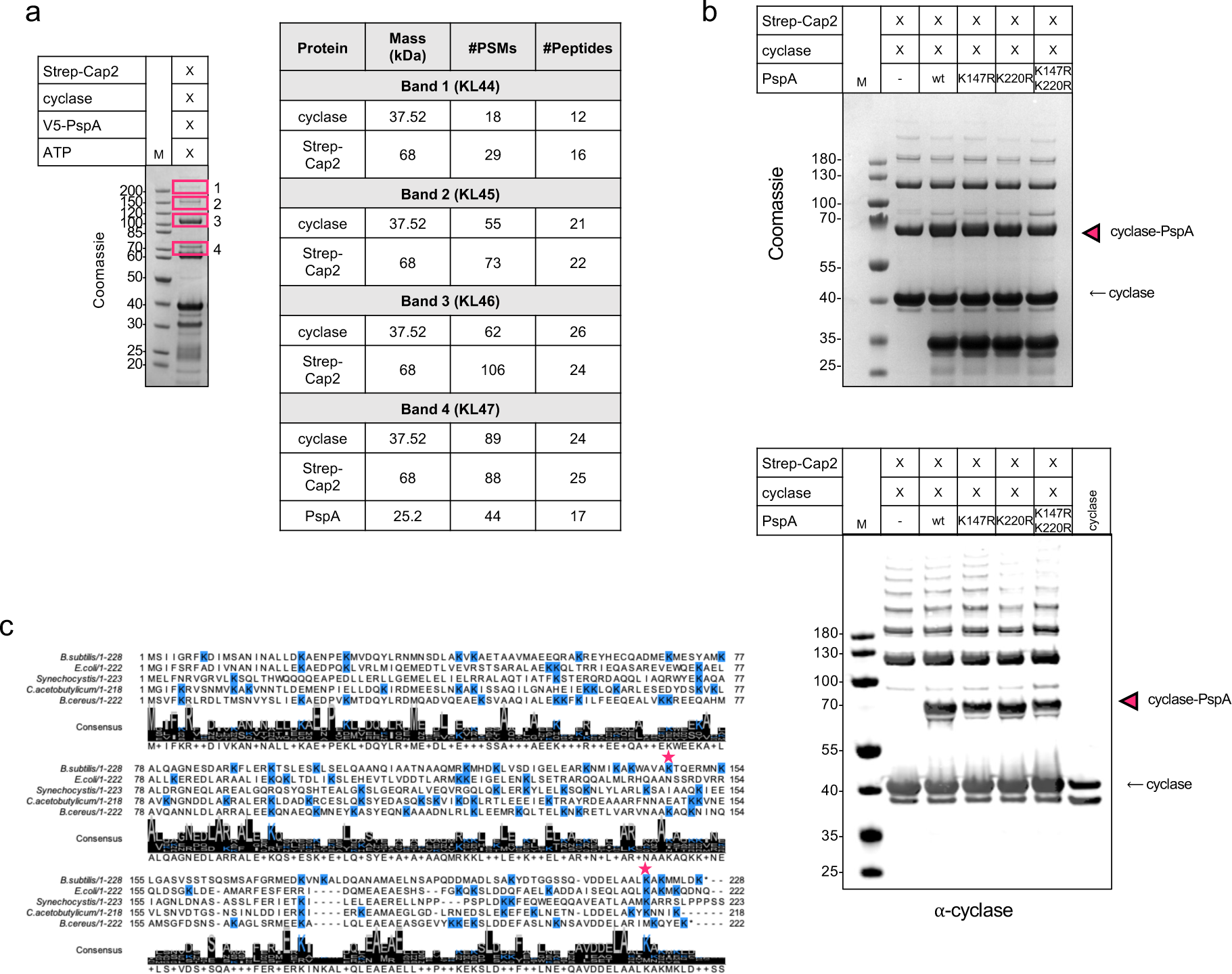
Cap2 conjugates the cyclase to PspA. **a**, *In vitro* conjugation of the cyclase to PspA. Reaction mixture contained 5 µM Strep-Cap2, 5 µM cyclase and 10 µM wild type PspA, 10 mM MgCl_2_, 1 mM ATP, 100 mM Tris-HCl pH 8, 150 mM NaCl. Samples were incubated for 30 min at 25°C, separated by SDS-PAGE. Bands were excised from the SDS gel, digested with trypsin and analyzed by mass spectrometry. The results of the peptide search are shown in the table. Abbreviations: PSMs: peptide sequence matches. **b**, *In vitro* conjugation of the cyclase to PspA. Reaction mixture contained 5 µM Strep-Cap2, 5 µM cyclase and 10 µM wild type PspA or mutants as indicated, 10 mM MgCl_2_, 1 mM ATP, 100 mM Tris-HCl pH 8, 150 mM NaCl. Samples were incubated for 30 min at 25°C, separated by SDS-PAGE (top) and analysed by Western blot (bottom) with a cyclase-specific antibody. Western blot shows the full blot image of the same blot from Figure 3b. Strep-Cap2: 67.8 kDa; cyclase: 37.8 kDa; PspA: 25.1 kDa. **c**, Sequence alignment of PspA homologs from *Bacillus subtilis* 168, *Bacillus cereus* WPySW2, *Escherichia coli*, *Clostridium acetobutylicum* and *Synechocystis sp. 1* was created with Geneious 2022.2.1 and visualized with Jalview 2.11.2.5. Lysine residues are highlighted in blue. The residues K147 and K220 of *B. subtilis* PspA are marked with magenta asterisks.

**Extended Data Figure 6.**
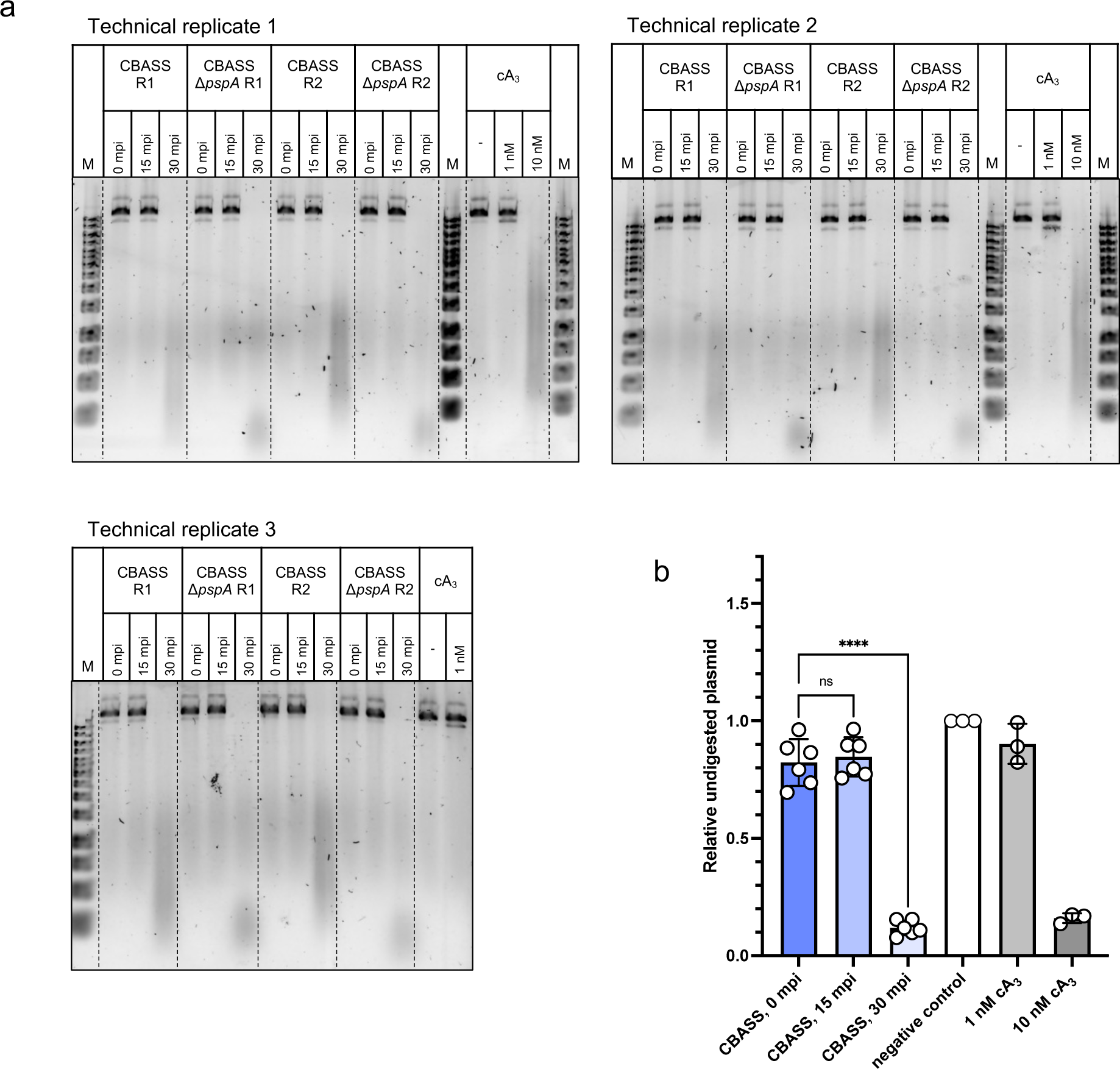
CBASS cyclic nucleotides can be detected in cells 30 min post infection. **a**, Nucleotides were extracted from *B. subtilis* cells carrying CBASS versions as indicated after infection with phage Goe23. Cultures were harvested 0, 15, or 30 minutes post infection (mpi). Nucleotides were tested for their ability to activate the CBASS effector Nuc-SAVED in a plasmid cleavage assay and plasmid cleavage assessed using a 1 % agarose gel. **Abbreviation: R1 and R2: biological replicate 1 and 2. b**, quantification of uncleaved plasmid after 30 min incubation of the nucleotide extracts with the Nuc-SAVED protein relative to the untreated control. The experiments were conducted with two biological replicates with three technical replicates each. Data are presented as mean values ± SD. Statistical analysis was performed using a one-way ANOVA, followed by Tukey’s multiple comparisons test (**P < 0.0052).

**Extended Data Figure 7.**
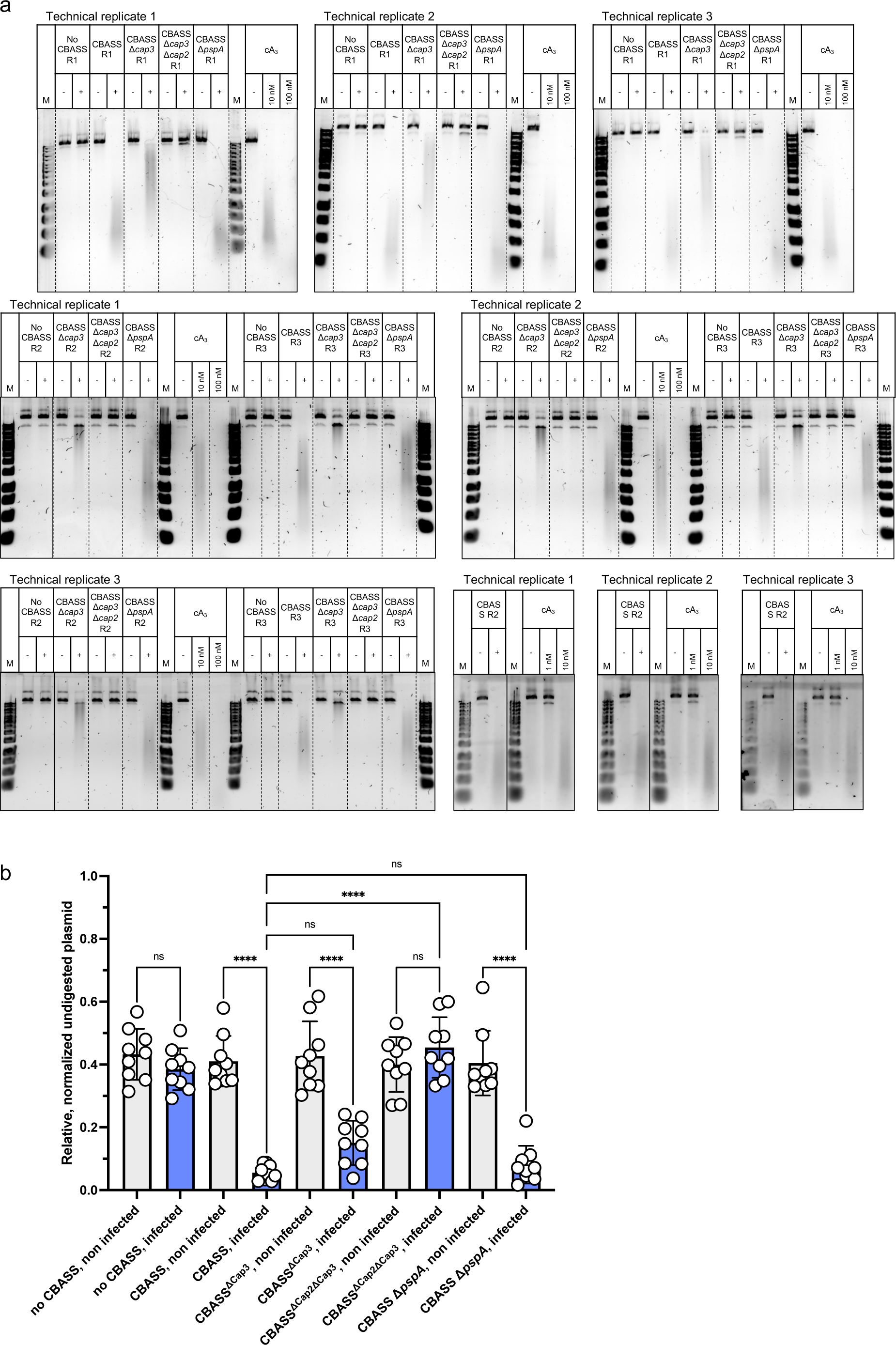
Production of CBASS cyclic nucleotides infected cells. **a**, *B. subtilis* wild type (no CBASS) and cells carrying CBASS versions as indicated were grown up to early exponential growth phase. Cultures were then either infected with the phage Goe23 (+) or not infected (-) and incubated for 30 min and harvested. Nucleotides were extracted and tested for their ability to activate the CBASS effector Nuc-SAVED in a plasmid cleavage assay. Plasmid cleavage was assessed using a 1 % agarose gel. **Abbreviation: R1 and R2: biological replicate 1 and 2. b**, quantification of uncleaved plasmid after incubation of the nucleotide extracts with the Nuc-SAVED protein relative to the untreated control and normalized to the protein content of the original cultures. The experiments were conducted with three biological replicates with three technical replicates each. Data are presented as mean values ± SD. Statistical analysis was performed using a one-way ANOVA, followed by Tukey’s multiple comparisons test (****P < 0.0001).

**Extended Data Figure 8.**
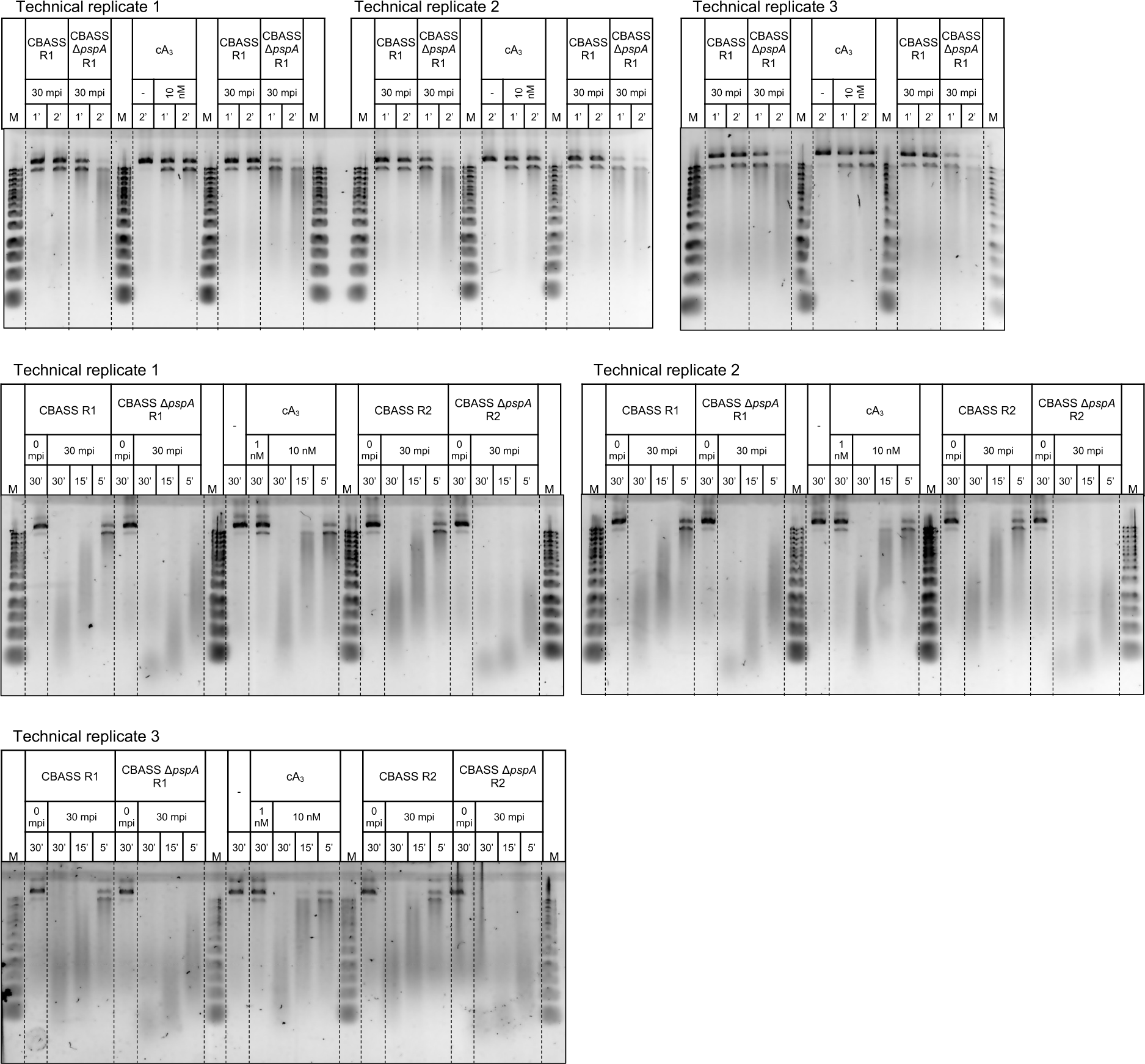
Increased second messenger production in the absence of PspA. Nucleotides were extracted from *B. subtilis* cells carrying CBASS versions as indicated before or 30 min after infection with phage Goe23. Nucleotides were tested for their ability to activate the CBASS effector Nuc-SAVED in a time course experiment (1, 2, 5, 15, 30 min) using a plasmid cleavage assay. The cleavage activity of Nuc-SAVED after incubation with the nucleotide extracts was assessed using a 1 % agarose gel. **Abbreviation: R1 and R2: biological replicate 1 and 2; mpi: minutes post infection.**

**Extended Data Figure 9.**
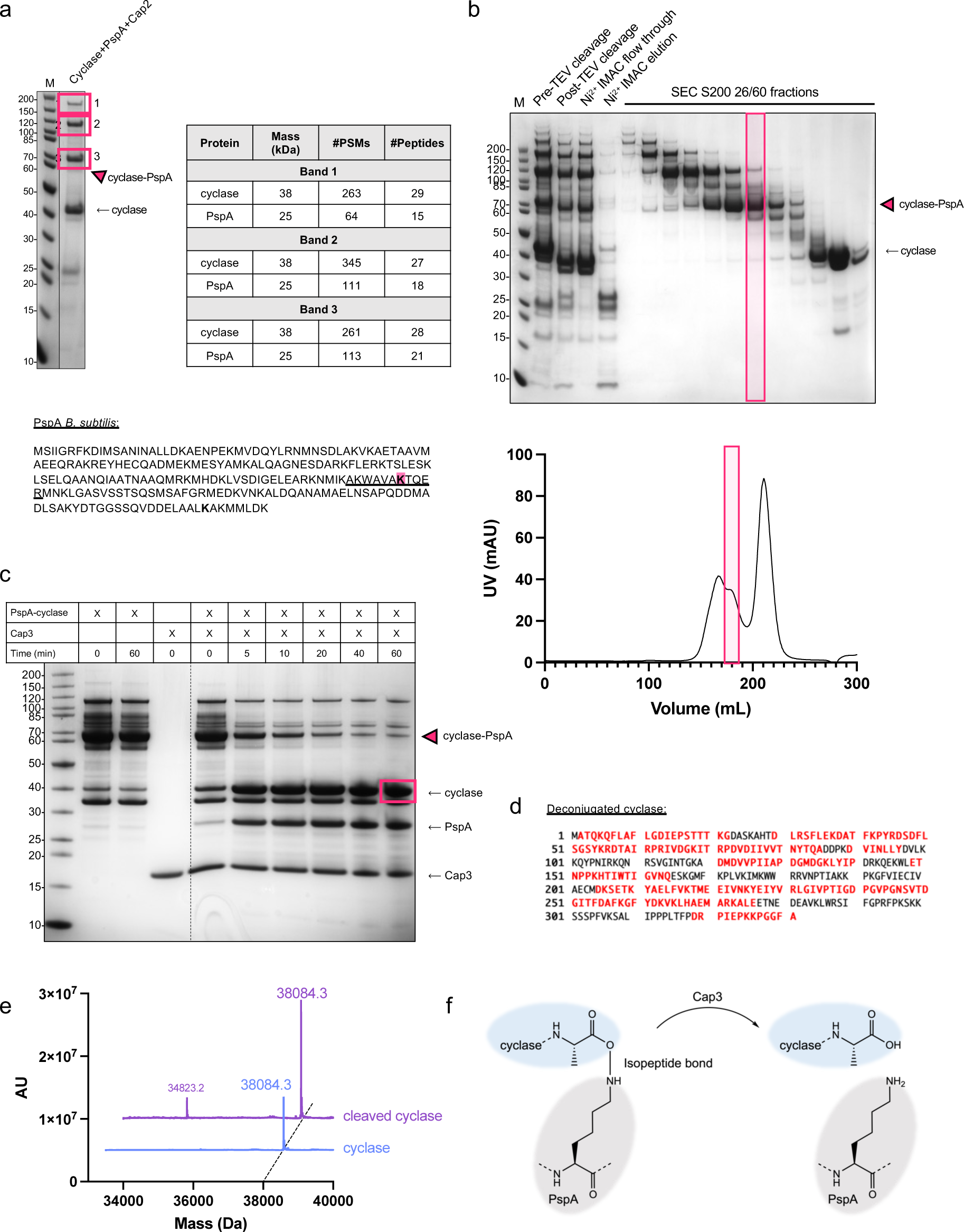
Conjugation in the heterologous *E. coli* host and cleavage of the conjugate by Cap3. **a**, The presence of cyclase and PspA peptides in the three conjugate bands was confirmed by mass spectrometry and the tryptic peptide (WAVAKTQER/AKWAVAKTQER) containing the modified lysine residues (highlighted in magenta) is emaild. The results of the peptide search are depicted in the table. Abbreviations: PSMs: peptide sequence matches. **b**, Co-expression of the CBASS cyclase (*Bce*), Cap2 (*Bce*) and PspA (*Bsu*) in *E. coli* and purification of the cyclase-PspA conjugate by size exclusion chromatography. **c**, the cyclase-PspA conjugate was incubated together with the Cap3 ortholog from *Cytobacillus oceanisediminis* and the cleavage was followed by SDS-PAGE (see also Fig. 5b). **d**, the band of the deconjugated cyclase was excised from the SDS gel (magenta box), digested with the peptidase AspN and the peptides were analysed by mass spectrometry. Peptides identified in the peptide search are shown in bold red. The overall protein sequence coverage was 61%. The presence of the C-terminal peptide including the C-terminal Ala residue involved in conjugation indicates that the cleavage of the cyclase-PspA conjugate by Cap3 leaves no scar on the cyclase. **e,** analysis of the size of the cyclase after cleavage. For this, the last sample (60 min) of the conjugate cleavage assay from c was analysed by intact mass spectrometry. **f**, cleavage of the cyclase-PspA conjugate by Cap3 leads to unaltered cyclase monomers.

**Extended Data Figure 10.**
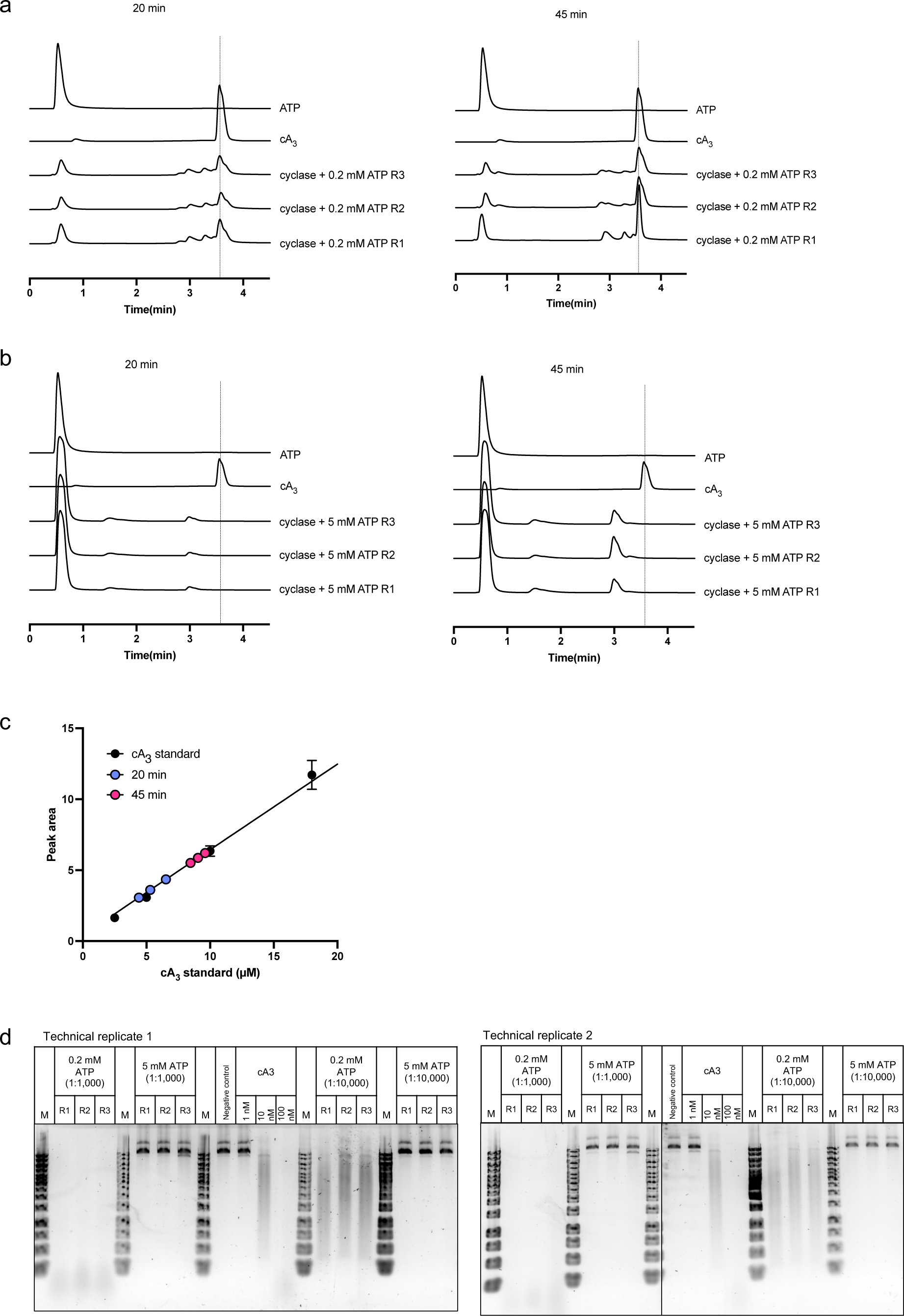
Analysis of cyclase products. **a**, reaction products from a cyclase reaction (50 µM cyclase, 0.2 mM ATP, 20 min / 45 min incubation at 37 °C) were analysed by HPLC. The traces of all three replicates are shown. Synthetic ATP and cA_3_ served as standard. **b**, same as **a** with the difference that the reaction mixture contained 5 mM ATP. **c**, a dilution series of synthetic cA_3_ was analysed by HPLC and the peak area of the cA_3_ peak used to create a standard curve. The amounts of cA_3_ in the reaction products from **a** were calculated by fitting to the standard curve. **d**, Nucleotides were tested for their ability to activate the CBASS effector Nuc-SAVED in a plasmid cleavage assay and plasmid cleavage assessed using a 1 % agarose gel.

